# Breakdown of sporophytic self-incompatibility: Diploids versus tetraploids

**DOI:** 10.64898/2026.07.27.740979

**Authors:** Diane Douet, Sylvain Billiard, Xavier Vekemans, Josselin Clo

## Abstract

Many angiosperm species possess self-incompatibility (SI) systems that prevent self-fertilization. Because empirical studies often report higher selfing rates in tetraploids than in diploids, we investigate whether sporophytic self-incompatibility (SSI) is more likely to break down after the introduction of a self-compatible (SC) allele in tetraploid populations. To address this question, we use analytical models and individual-based simulations to compare diploid and tetraploid populations under two main scenarios: (1) all SI alleles are codominant, and (2) SI alleles are structured into dominance classes. Overall, our results indicate that SSI is more difficult to maintain in tetraploids than in diploids, with dominance relationships playing a key role in the invasion success of an SC allele. When SI alleles are organized into dominance classes, increasing the dominance of the SC allele generally favors SSI breakdown in tetraploids, while diploids show weaker sensitivity to dominance, with SSI maintained across all dominance scenarios for the SC allele under sufficiently high inbreeding depression. However, when the SC allele is dominant over all SI alleles, SSI is more readily maintained in both the codominant and dominance-class models.

## 1 Introduction

Polyploidy, characterized by the presence of more than two sets of homologous chromosomes, is especially widespread in plants, as around 35% of angiosperms are recent polyploids (Wood et al., 2009; Heslop-Harrison et al., 2023), and many angiosperm families show evidence of ancient genome duplications (Jiao et al., 2011; Van de Peer et al., 2017). Polyploidy is also thought to facilitate speciation and adaptation (Otto and Whitton, 2000; Van de Peer et al., 2021). Despite its prevalence, the establishment of neo-polyploids in diploid populations is hindered by several constraints. First, whole-genome duplication can induce genomic instability, mitotic and meiotic abnormalities, and reduced fitness (Comai, 2005; Otto, 2007; Doyle and Coate, 2019; Porturas et al., 2019; Clo and Kolář, 2021). In addition, neo-polyploids are subject to a frequency-dependent disadvantage in which rare cytotypes experience reduced reproductive success called “minority cytotype exclusion” (MCE) (Levin, 1975; Ramsey and Schemske, 1998). For this reason, the way that polyploids arise and establish in diploid populations remains unclear and has raised particular interest over the last decades (see e.g. Osterman et al. (2025)), with many models exploring hypotheses such as a fitness advantage (Burton and Husband, 2000; Porturas et al., 2019; Clo and Kolář, 2021; Cheng et al., 2026), an elevated rate of unreduced gamete production (Felber, 1991; Clo et al., 2022; Gerstner et al., 2024), or reproductive isolation (Baack, 2005; Griswold, 2021; Kauai et al., 2025; Zwaenepoel, 2025) to explain it.

Self-fertilization has long been proposed as a mechanism that can facilitate polyploid establishment because it can reduce the impact of inter-cytotype mating. By providing reproductive assurance and reducing gene flow from surrounding diploid populations, increased selfing can help newly formed polyploids overcome MCE (Antonovics, 1968; Jain, 1976; Rausch and Morgan, 2005). Empirical studies also report frequent associations between polyploidy and self-fertilization (Stebbins, 1950; Barringer, 2007). Moreover, the evolution of selfing in polyploids may be facilitated by reduced inbreeding depression compared to their diploid counterparts, particularly during early stages of establishment (Clo and Kolář, 2022; Orsucci et al., 2022). However, polyploidy may instead lead to higher inbreeding depression under certain genetic assumptions, such as changes in the dominance of deleterious alleles under tetrasomic inheritance (Ronfort, 1999). The relationship between polyploidy and selfing remains debated (Mable, 2004) and seems to vary between autopolyploids and allopolyploids (Husband et al., 2008; Vekemans et al., 2026), highlighting the complexity of the relationship between polyploidy and the evolution of self-fertilization. Moreover, self-fertilization is often prevented by self-incompatibility (SI) systems, which occur in about 45% of angiosperm species (Ferrer et al., 2025). Increased selfing rates in polyploids therefore require the breakdown of SI systems when they remain functional, causing a transition from self-incompatibility (SI) to self-compatibility (SC), which has been documented in several empirical (Orsucci et al., 2022; Kolesnikova et al., 2023; Novikova et al., 2023) or experimental studies (Yew et al., 2023; Duan et al., 2024). More theoretical investigations are therefore necessary to understand whether the breakdown of SI systems is facilitated in tetraploids compared to diploids and under which conditions the loss of SI occurs.

SI systems are generally classified into two types: gametophytic SI (GSI), where the pollen phenotype is determined by its haploid genotype, and sporophytic SI (SSI), where it is determined by the diploid genotype of the parent plant (Bateman, 1952). In GSI systems, polyploidization can weaken self-incompatibility and promote selfing (Stone, 2002), and in GSI with nonself-recognition, it may even directly cause the transition to self-compatibility (Fujii et al., 2016), facilitating tetraploid establishment under certain conditions (Douet et al., 2026). By contrast, the breakdown of GSI and SSI systems with self-recognition in polyploids necessarily involves the introduction of a non-functional self-compatible S-allele, or an unlinked modifier allele (Duan et al., 2024). However, the fate of a self-compatible allele in SSI systems has received little attention in theoretical population genetics. SSI is particularly common in the *Brassicaceae* family, where it relies on the interaction between the S receptor kinase (*SRK* ) in the pistil and the S-locus cysteine-rich (*SCR/SP11* ) protein in the pollen (Schopfer et al., 1999; Takasaki et al., 2000). Loss-of-function mutations in either the *SRK* or the *SCR* genes can disrupt SI systems (Goring et al., 1993; Tsuchimatsu et al., 2010), producing non-functional S-alleles that allow self-fertilization depending on the dominance of the parental S-allele. Indeed, although codominance is mostly observed in the pistil (Hatakeyama et al., 2001; Prigoda et al., 2005), dominance relationships in the pollen generally result in the expression of a single S-allele in heterozygous individuals in *Brassicaceae* (Schoen and Busch, 2009; Fujii and Takayama, 2018). Dominance appears to be a key factor influencing the outcome of SSI systems in polyploids (Vekemans et al., 2026; Yew et al., 2023). In particular, it can contribute to SI breakdown or maintenance in hybrids depending on interactions among S-alleles (Duan et al., 2024), and it can determine the transition to self-compatibility in allopolyploids through interactions between parental S-haplotypes (Novikova et al., 2023). Theoretical studies on the transition from SI to SC have identified several key factors.

In particular, the role of inbreeding depression and the degree of severity of deleterious mutations have been investigated extensively. These studies show that alleles increasing selfing rates can be favored by selection and spread in populations even under high levels of inbreeding depression, partly due to increased homozygosity that facilitates the purging of highly deleterious mutations (Charlesworth et al., 1990; Lande et al., 1994). However, in small populations, higher selfing rates can accelerate the accumulation of mildly deleterious mutations and increase the probability of extinction (Abu Awad and Billiard, 2017). Moreover, pollen discounting can also significantly reduce the benefit of selfing as selfing rates increase (Holsinger, 1991; Johnston, 1998). As a result, SI breakdown depends on a balance between the automatic advantage of selfing, reproductive assurance, and inbreeding depression (Gervais et al., 2014). Another factor that has been identified to promote a transition from SI to SC is pollen limitation. Theoretical studies have shown that pollen-limited environments can favor SC (Charlesworth and Charlesworth, 1979; Goodwillie, 1999; Porcher and Lande, 2005a), as SI becomes disadvantageous due to an Allee effect that reduces S-locus diversity and therefore the number of compatible mates (Wagenius et al., 2007; Leducq et al., 2010). However, most of these models have been developed for GSI systems in diploid populations, and they remain to be extended to SSI systems, as well as to polyploid populations, to improve our understanding of the evolution of mating strategies in flowering plants.

In this article, we investigate the outcome of SSI systems following the introduction of a mutant SC allele in diploid and tetraploid populations, focusing on the effects of increased self-pollen rates and inbreeding depression. By comparing diploid and tetraploid populations, we aim to determine whether the breakdown of SSI is more likely in tetraploids than in diploids, and how dominance relationships affect the invasion and maintenance of self-compatible alleles. Because the loss of SSI may facilitate the evolution of selfing in newly formed polyploids, our results may also provide insights into one potential mechanism contributing to neo-polyploid establishment. To address this question, we use analytical modeling and individual-based simulations, extending the framework of Gervais et al. (2014) to SSI systems and tetraploids. We consider two dominance scenarios for S-alleles (Figure 1): (1) all SI alleles are codominant, and (2) SI alleles are structured into dominance classes within which alleles are codominant. The different scenarios are used to explore the influence of dominance relationships among different SI alleles and dominance between SI alleles and the SC mutant on the maintenance or breakdown of SSI at both ploidy levels. Figure 1 shows that the ploidy level and dominance relationships can strongly impact SSI systems. Our study is motivated by these effects, inducing the possibility of various outcomes for the invasion conditions of a mutant self-compatible allele.

**Figure 1:**
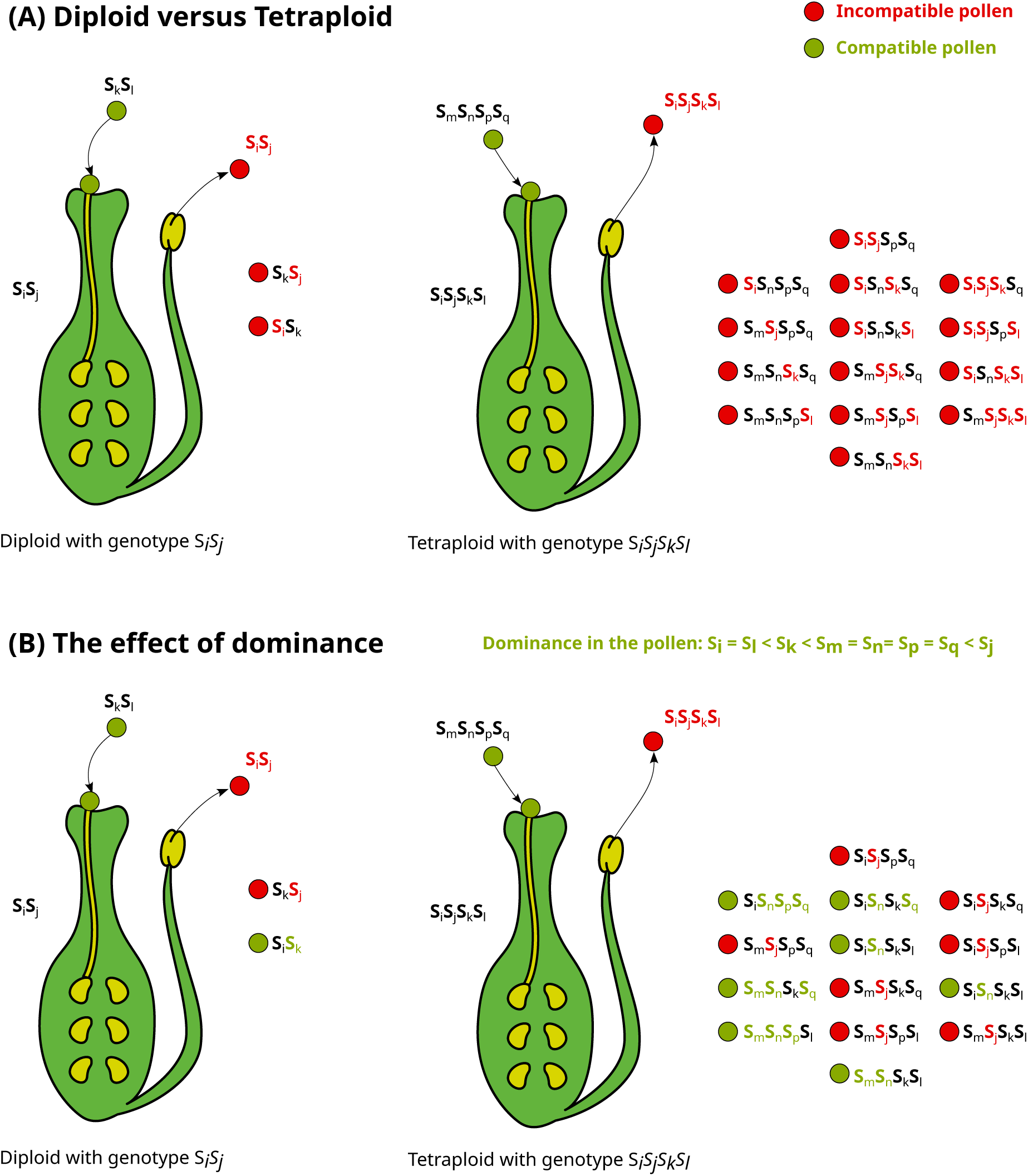
The effects of ploidy (with S-alleles under strict codominance) (A) and dominance relationships (B) on the number of compatible mates in sporophytic self-incompatibility. Polyploidy makes SSI systems more restrictive (A), while dominance effects between SI alleles in the pollen can increase the pool of compatible mates at both ploidy levels (B).

## 2 Models and methods

Throughout this paper, an SC allele refers to a non-functional S-allele resulting from a loss-of-function mutation at the S-locus, whereas an SI allele refers to a functional S-allele with intact pollen and pistil component genes. Note that the presence of an SC allele in a genotype does not automatically confer self-compatibility. The self-compatible (SC) or self-incompatible (SI) phenotype of an individual depends on the number of SC alleles it carries and on the dominance relationships between SC and SI alleles. The notations and parameter values used in the models and simulations are detailed in Table 1. We developed models covering all combinations of SI dominance relationships, ploidy levels, modeling approaches, and SC allele dominance scenarios presented in Table 2, unless otherwise stated.

**Table 1:** Summary of notations and parameter values.

| Symbol | Meaning | Default values |
| --- | --- | --- |
| $N_{\text{pop}}$ | Population size | 1000 |
| $W_i$ | Fitness of individual $i$ | |
| $s$ | Selection coefficient of deleterious mutations | 0.05 |
| $n$ | Number of SI alleles in the population | |
| $h$ | Dominance coefficient of deleterious mutations in diploids | 0.25 |
| $h_1, h_2, h_3$ | Dominance coefficients of deleterious mutations in tetraploids | 0.1, 0.2, 0.3 |
| $\alpha$ | Self-pollen rate | |
| $\delta$ | Inbreeding depression | |
| $L$ | Chromosome map length | 10 |
| $U$ | Mutation rate per haplotype | |
| $U_{SI}$ | Mutation rate at the S-locus from SI to SI | $10^{-5}$ |
| $U_{SC}$ | Mutation rate at the S-locus from SI to SC | $10^{-4}$ |

**Table 2:** Summary of the models developed. IBM stands for individual-based model.

| SI dominance | Ploidy | Model | $S_C$ dominance |
| --- | --- | --- | --- |
| Codominant SI alleles (Section 2.1). | • Diploids<br>• Tetraploids | • IBM: fixed inbreeding depression<br>• IBM: deleterious mutations<br>• Analytical model | • Codominant/recessive to all SI alleles.<br>• Dominant over all SI alleles. |
| SI alleles in dominance classes (Section 2.2). | • Diploids<br>• Tetraploids | • IBM: fixed inbreeding depression<br>• Deterministic model (diploids only) | • Belongs to one of the classes 1-4.<br>• Dominant over all SI alleles (class 5). |

### 2.1 Codominant SI alleles

In this section, all self-incompatible (SI) S-alleles are considered codominant (which corresponds to the model SSIcod in Schierup et al. (1997)). The population is modeled under a sporophytic self-incompatibility system (SSI) with self-recognition. Both SC and SI alleles are assumed to be codominant in the pistil. In the pollen, all SI alleles are assumed to be codominant, and two scenarios are considered for the dominance relationships of the SC allele: either the SC allele is dominant over all SI alleles, or it is codominant with or recessive to the SI alleles. Note that when *S_C_* is dominant over all SI alleles, any individual carrying at least one SC allele is self-compatible, whereas when *S_C_*is codominant or recessive, only homozygous *S_C_S_C_* individuals (or *S_C_S_C_S_C_S_C_*in tetraploids) are self-compatible, therefore determining the dominance of the self-compatible phenotype. The models developed in this section are extensions of the models presented in Gervais et al. (2014), in which they consider diploid populations under gametophytic self-incompatibility.

#### 2.1.1 Analytical model: Diploids

We develop a deterministic model in which our population of mutants, i.e., individuals in the population that have at least one *S_C_* allele, is structured in different groups. We denote by *x*_1_ and *x*_2_ the frequencies of individuals with genotype *S_C_S_C_* produced by selfing and outcrossing, respectively. Similarly, we denote by *x*_3_ and *x*_4_ the frequencies of *S_i_S_C_* individuals produced by selfing and outcrossing, respectively, where *S_i_* denotes any SI allele. We also denote

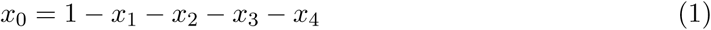

the frequency of *S_i_S_j_*individuals. We assume that selfed individuals experience inbreeding depression, such that *W*_0_ = *W*_2_ = *W*_4_ = 1, while *W*_1_ = *W*_3_ = 1 − *δ*. The mean fecundity is therefore expressed by 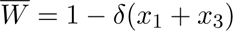.

Various dominance relationships in the pollen between the *S_C_* allele and all the other SI alleles are considered:

- *S_C_* **codominant or recessive**: In that case, only *S_C_S_C_* individuals are self-compatible. The recursion equations are available in Section A.1.1 of the appendix.
- *S_C_* **dominant**: In that case, both *S_C_S_C_* and *S_C_S_i_*individuals are self-compatible for all *i*. Note that *S_i_S_i_*individuals produced via selfing can appear in the population and suffer from inbreeding depression. This fitness difference is not considered here, as analyses have shown that including or neglecting this fitness difference leads to similar outcomes (see *Wolfram Mathematica* Notebook for more details and Figure S1 of the appendix for comparison). The recursion equations are available in Section A.1.1 of the appendix.

The selfing rates for self-compatible groups are denoted by *a*_1_, *a*_2_, *a*_3_ and *a*_4_, respectively, and are computed as follows:

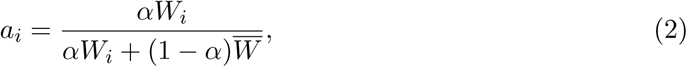

where *α* is the self-pollen rate, i.e. the proportion of pollen that remains on the maternal plant and is available for self-fertilization. For self-incompatible groups, *a_i_* = 0.

In both scenarios, the frequencies *x*_1_*^′^ , x*_2_*^′^ , x*_3_*^′^* and *x*_4_*^′^* at the next generation are obtained by summing the frequencies of all reproductive events (through outcrossing or selfing, depending on the group) that generate individuals of the corresponding group. Section A.2.1 of the appendix provides a detailed analytical resolution of both models using Routh-Hurwitz stability conditions after linearization to first order.

#### 2.1.2 Analytical model: Tetraploids

We structure our population of mutants, i.e., individuals in the population that have at least one *S_C_* allele, into different groups. We denote by *x*_1_*_s_*, *x*_2_*_s_*, *x*_3_*_s_* and *x*_4_*_s_* the frequencies of individuals with 1,2,3 or 4 *S_C_*alleles produced by selfing. Similarly, we denote by *x*_1_*_c_*, *x*_2_*_c_*, *x*_3_*_c_* and *x*_4_*_c_* the frequencies of individuals with 1,2,3 or 4 *S_C_*alleles produced by outcrossing. We also denote:

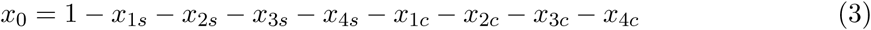

the frequency of *S_i_S_j_S_k_S_l_* individuals (with no *S_C_* alleles). We assume that selfed individuals experience inbreeding depression, such that *W*_0_ = *W*_1_*_c_* = *W*_2_*_c_* = *W*_3_*_c_* = *W*_4_*_c_* = 1, while *W*_1_*_s_* = *W*_2_*_s_* = *W*_3_*_s_* = *W*_4_*_s_* = 1 − *δ*. The mean fecundity is therefore expressed by 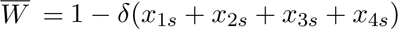.

Various dominance relationships in the pollen between the *S_C_* allele and all the other *S_i_* alleles are considered:

- *S_C_* **codominant or recessive**: In that case, only fully homozygous individuals at the S-locus (*S_C_S_C_S_C_S_C_*individuals) are self-compatible. The recursion equations are available in Section A.1.2. Section A.2.2 of the appendix provides a detailed analytical resolution of this model for tetraploids using Routh-Hurwitz stability conditions after linearization to first order.
- *S_C_* **dominant**: In that case, all individuals that have at least one *S_C_* allele are self-compatible, and their pollen grains are compatible with all individuals in the population. Similarly to diploids, individuals carrying four SI alleles can arise in the population through selfing when the SC allele is dominant. Since the probability of producing such individuals is comparable between diploid and tetraploid populations, we neglect this fitness difference here. The recursion equations are available in Section A.1.2. This model is solved numerically by iterating the recursion equations until an equilibrium is reached.

The selfing rates for self-compatible groups are denoted by *a*_1*s*_, *a*_2*s*_, *a*_3*s*_ and *a*_4*s*_ for selfed individuals, and *a*_1*c*_, *a*_2*c*_, *a*_3*c*_ and *a*_4*c*_ for outcrossed individuals, and are computed as follows:

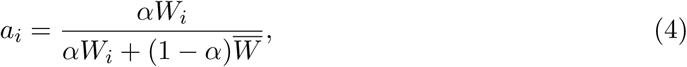

where *α* is the self-pollen rate. For self-incompatible groups, *a_i_* = 0.

#### 2.1.3 Individual-based simulations

A simulation model is developed to determine the level of inbreeding depression required to prevent the invasion of an SC allele in a population of *N*_pop_ diploid or tetraploid individuals. At the beginning of the simulation, *N*_pop_ diploid (resp. tetraploid) individuals are created, each carrying two (resp. four) chromosomes represented by empty lists that will further store the position of deleterious mutations. Two (resp. four) S-alleles, located at the center of the chromosomes, are randomly assigned to each individual from a pool of *k* = 100 possible S-alleles.

The next generation is obtained through a life-cycle consisting of selection, recombination, reproduction, and mutations. For each offspring, a maternal parent is first selected with probability proportional to its fitness *W_i_*. When the value of inbreeding depression is fixed throughout the simulation, the fitness of an individual *i* is defined as:

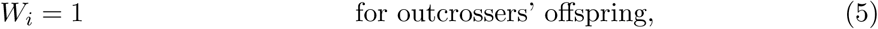

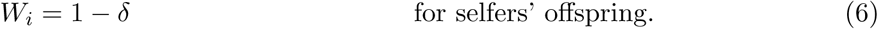

When inbreeding depression is controlled by the deleterious mutation rate *U* , the fitness of an individual is defined as:

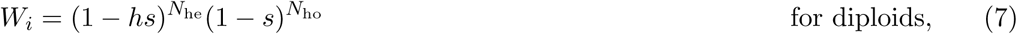

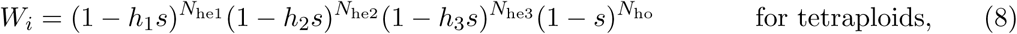

where *h*, *h*_1_, *h*_2_ and *h*_3_ denote the dominance coefficients associated with heterozygous genotypes and *s* is the selection coefficient of deleterious mutations. *N*_he_, *N*_he1_, *N*_he2_, and *N*_he3_ are the numbers of heterozygous mutations (on 1, 2, or 3 chromosomes, respectively, for tetraploids) and *N*_ho_ is the number of homozygous mutations carried by the individual. The maternal parent self-fertilizes with a probability equal to its selfing rate *a_i_*, computed as follows:

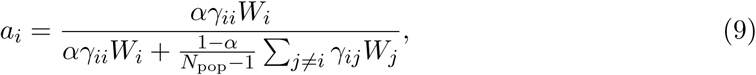

where *α* is the self-pollen rate and *γ_ij_*is the number of gametes produced by individual *j* that are compatible with individual *i*.

If self-fertilization occurs, an offspring is created from two (for diploids) or four (for tetraploids) recombined chromosomes of the maternal parent and added to the next generation. Note that all pollen grains produced by an SC individual are compatible with its own pistil in that case.

If the maternal parent engages in outcrossing, a paternal parent is selected with probability proportional to its fitness *W_j_*, and its compatibility with the mother at the S-locus is evaluated. When the SC allele is dominant over SI alleles, the pollen is compatible if at least one S-allele carried by the paternal parent is SC, or if all of its S-alleles are different from those of the maternal parent. When the SC allele is codominant with or recessive to the SI S-alleles, the pollen is compatible only if all paternal S-alleles are SC, or if all of its SI S-alleles are different from those of the maternal parent. In the diploid case, if the selected paternal parent is incompatible, another candidate is drawn from the population until a compatible father is found, therefore representing an environment without pollen limitation. In tetraploids, the increased number of S-allele copies reinforces self-incompatibility, making compatible mates harder to find. In that case, another maternal parent is selected if compatible pollen is not found after 50 attempts to ensure smooth running of the simulation while still considering low pollen limitation. Additional simulations with higher pollen limitation (5 attempts maximum before rejecting the mother) confirmed that pollen limitation does not have a significant impact on the results, as presented in Figure S2.

Once compatible parents are selected, gametes are formed by recombination. The number of crossovers is drawn from a Poisson distribution with parameter *L*, and their positions are uniformly sampled between 0 and 1. Offspring genomes are then subjected to various types of mutations depending on the phase of the simulation.

A simulation run consists of three phases in which different types of mutation occur. In the first phase, which lasts 2000 generations, each SI allele can mutate to another SI allele among the *k* = 100 possible alleles, with probability *U_SI_*. The number of such mutations is drawn from a binomial distribution with parameters (ploidy × *N*_pop_*, U_SI_*). For each mutation, an individual and one of its S-alleles are chosen uniformly. In the second phase, lasting 2000 generations, deleterious mutations are also introduced. This phase is unnecessary when inbreeding depression is fixed. For each chromosome, the number of deleterious mutations is drawn from a Poisson distribution with parameter *U* , and their positions on the chromosome are uniformly sampled between 0 and 1, corresponding to an infinite number of loci. In the third and final phase, S-alleles can mutate to the SC allele with probability *U_SC_*, in addition to the other types of mutations. The number of such mutations is drawn from a binomial distribution with parameters (ploidy × *N*_pop_*, U_SC_*), after which each mutation is assigned to a uniformly chosen individual and one of its S-alleles. This phase lasts for up to 250, 000 generations, but stops earlier (after at least 50, 000 generations) if the SC allele invades the population, as no reverse mutation from SC to SI is allowed.

At the end of each simulation, the frequency of SC alleles in the population is computed. The number of S-alleles present in the population and the level of inbreeding depression are computed at the end of the second phase and also returned at the end of the simulation.

To determine the minimal level of inbreeding depression required to prevent invasion of the SC allele, we use a dichotomous search, running multiple simulations. The mutation rate *U* is increased when the frequency of the SC allele exceeds 0.05 at equilibrium, and decreased otherwise. To target a specific value of the inbreeding depression *δ*, a dichotomy is performed on the first two phases of the simulation until the difference between the observed and desired values inbreeding depression *δ* is less than 10*^−^*^2^, therefore calibrating *U* before introducing SC alleles in the final phase.

### 2.2 SI alleles structured in dominance classes

In this section, SI alleles are assumed to be always codominant in the pistil, but a combination of codominant and dominant interactions are occurring in the pollen (which corresponds to the domcod model in Billiard et al. (2007)). In the pollen, SI alleles are structured in four dominance classes, with codominance among alleles within classes and dominance between classes. Class 1 represents the most recessive alleles and class 4 the most dominant ones. The number of dominance classes was chosen based on empirical studies in the Brassicaceae family, particularly in outcrossing *Arabidopsis* species, which have documented the presence of four dominance classes in pollen (Prigoda et al., 2005; Durand et al., 2014; Vekemans et al., 2026). Several scenarios are considered for the dominance relationships of the SC allele in the pollen: either the SC allele belongs to one of the dominance classes 1-4, or it is dominant over all SI alleles (class 5).

#### 2.2.1 Analytical model: Diploids

To see the impact of dominance on the loss of SSI, we developed a deterministic model assuming that SI alleles belong to different dominance classes, and that the SC allele either belongs to one of those classes or is dominant over all SI alleles. In this case, we compute the frequency of all genotypes *S_i_S_j_*, where *i, j* ∈ [0*, n* ] with *n* the number of SI alleles in the population and *S*_0_ represents *S_C_*. We structure our population in two categories: the genotypes produced by selfing with frequency *x^s^* and those produced by outcrossing with frequency *x^c^* , as we assume that individuals produced by selfing will have a lower fitness *W^s^* = 1 − *δ* compared to those produced by outcrossing *W ^c^* = 1.

The selfing rate of a self-compatible individual with genotype *S_i_S_j_*is given by the following expression:

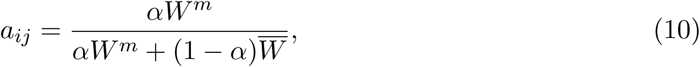

where *W^m^* = *W^c^* for individuals produced by outcrossing, and *W^m^* = *W^s^* for individuals produced by selfing. If the individual is self-incompatible, i.e. if the most dominant S-allele among *S_i_* and *S_j_* is an SI allele, *a_ij_* = 0. The recursion equations are available in Section A.1.3 of the appendix.

This model is solved numerically, by implementing functions in *Julia* that compute the compatibility between two individuals, their probability of producing a given offspring, and their frequency at the next generation. The population is represented as a 3-dimensional matrix of size (*n* + 1) × (*n* + 1) × *k*, where dimension *k* = 2 is to distinguish between offspring from outcrossing versus selfing. The number of SI alleles *n* is fixed and given by the results from individual-based simulations, as well as the distribution of those alleles into the different classes.

#### 2.2.2 Individual-based model

We modified the individual-based model presented in Section 2.1.3 to perform simulations in which we consider four dominance classes for S-alleles, with class 1 being the most recessive and class 4 being the most dominant. The SC allele either belongs to one of these classes or belongs to class 5 (dominant over all SI alleles). In that case, the S-alleles are still assumed to be codominant in the pistil. In this simulation, *S_i_* belongs to class 1 if *i* ∈ [1, 25]], to class 2 if *i* ∈ [26, 50]], to class 3 if *i* ∈ [51, 75]], and to class 4 if *i* ∈ [76, 100]]. Pollen is compatible only if the most dominant paternal S-allele is SC, or if the most dominant of both SI S-alleles is different from those of the maternal parent.

For this section, we fixed the value of inbreeding depression for all simulations instead of considering deleterious mutations to avoid any numerical issues. To find the inbreeding depression threshold over which the SC allele is absent from the population, we used a dichotomous search, stopping the simulation when the difference between *δ*_max_ and *δ*_min_ is less than 10*^−^*^2^. The SC allele is considered to be absent from the population when the frequency of the SC allele is lower than 0.05 after 250, 000 generations.

## 3 Results

### 3.1 Codominant SI alleles

We consider the scenario where all SI alleles are codominant and investigate the impact of the dominance of the SC allele, as well as the impact of the ploidy level, on the maintenance or breakdown of the SSI system. This analysis is based both on the predictions of the analytical model and on the results obtained using individual-based simulations described in Section 2.1.

#### In diploids

When the SC allele is dominant over all SI alleles, Figure 2 shows that for high levels of inbreeding depression, the SC allele fails to be maintained in the population when the self-pollen rate exceeds 0.2. In this case, simulation results closely match analytical predictions, as inbreeding depression is treated as a fixed parameter in the individual-based model. By contrast, larger quantitative differences between simulation and analytical results are observed in Figure S3 where inbreeding depression can fluctuate throughout the simulations, as it is determined by the rate of deleterious mutations *U* . Introducing deleterious mutations also leads to variation in the number of SI alleles maintained in the population before the introduction of the SC allele, which further impacts its maintenance in the population. However, as these discrepancies remain limited, treating inbreeding depression as a fixed parameter represents a reasonable approximation, and we therefore use this approach in the following simulations.

**Figure 2:**
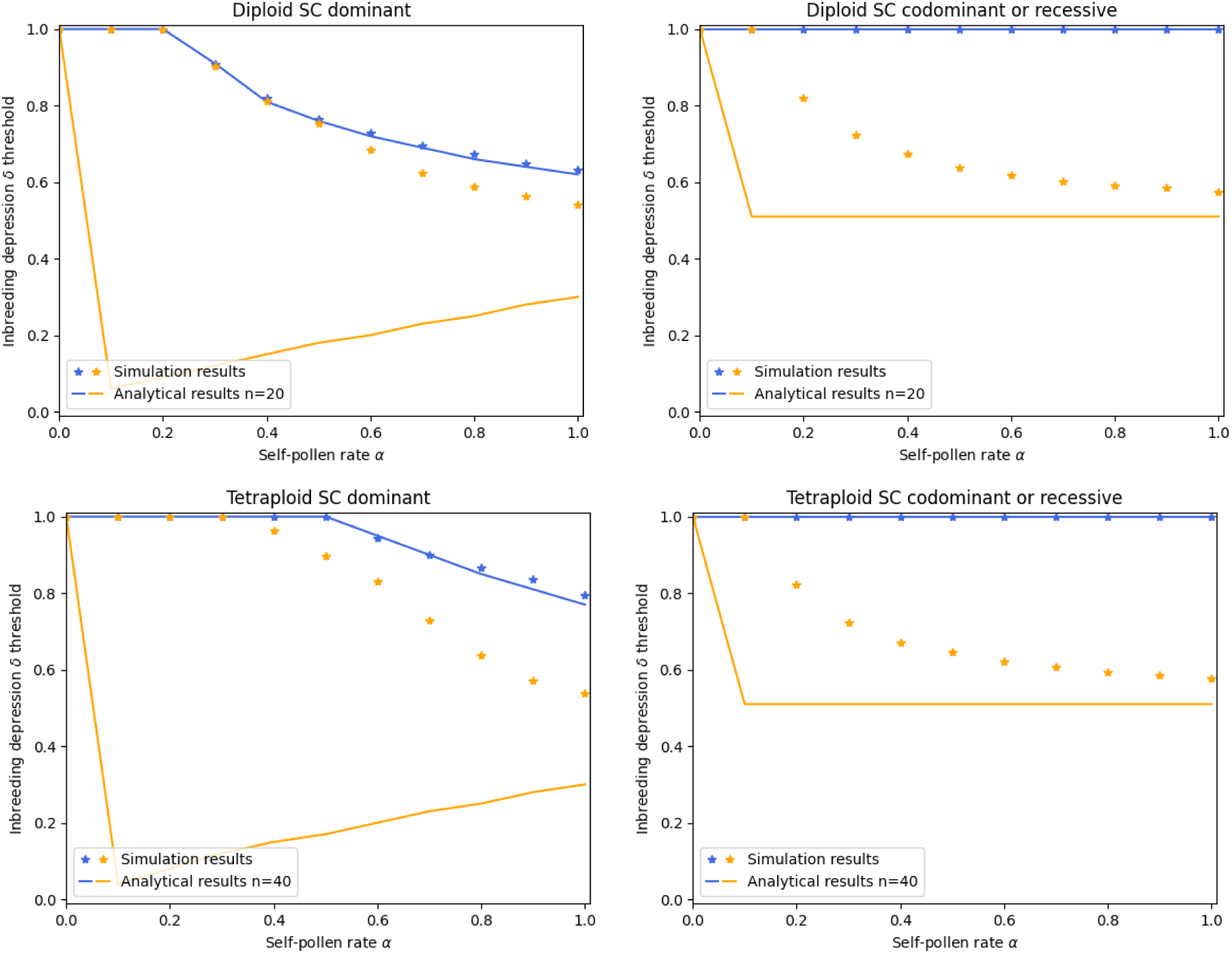
SI codominant. Blue curves represent the minimal inbreeding depression necessary to prevent the maintenance of an SC mutant allele into an initially SI population of *N*_pop_ = 1000 individuals. Orange curves represent the inbreeding depression threshold from which coexistence between SI and SC alleles occurs. More specifically, the SSI system is fully maintained for parameters above the blue curve, while the SC allele is present in the population at equilibrium for parameters below the curve, coexisting with SI alleles between the two curves, and fixed in the population below the orange one. Comparison between results obtained using individual-based simulations with ten replicates (star symbol) where inbreeding depression is fixed and analytical results (solid lines) for a diploid (top) and a tetraploid population (bottom), when the SC allele is dominant over all SI alleles (left), or recessive/codominant (right). The numbers of SI alleles in the analytical model (*n* = 20 in diploids and *n* = 40 in tetraploids) were chosen to reflect the values observed in simulations before introducing the SC allele.

In contrast, when the SC allele is recessive or codominant, both approaches reveal that the SC allele is always present at equilibrium in the population regardless of the level of inbreeding depression, and this for all selfing rates (see Figures 2, 3 and 4, and Section A.2.1 of the appendix for the analytical resolution).

**Figure 3:**
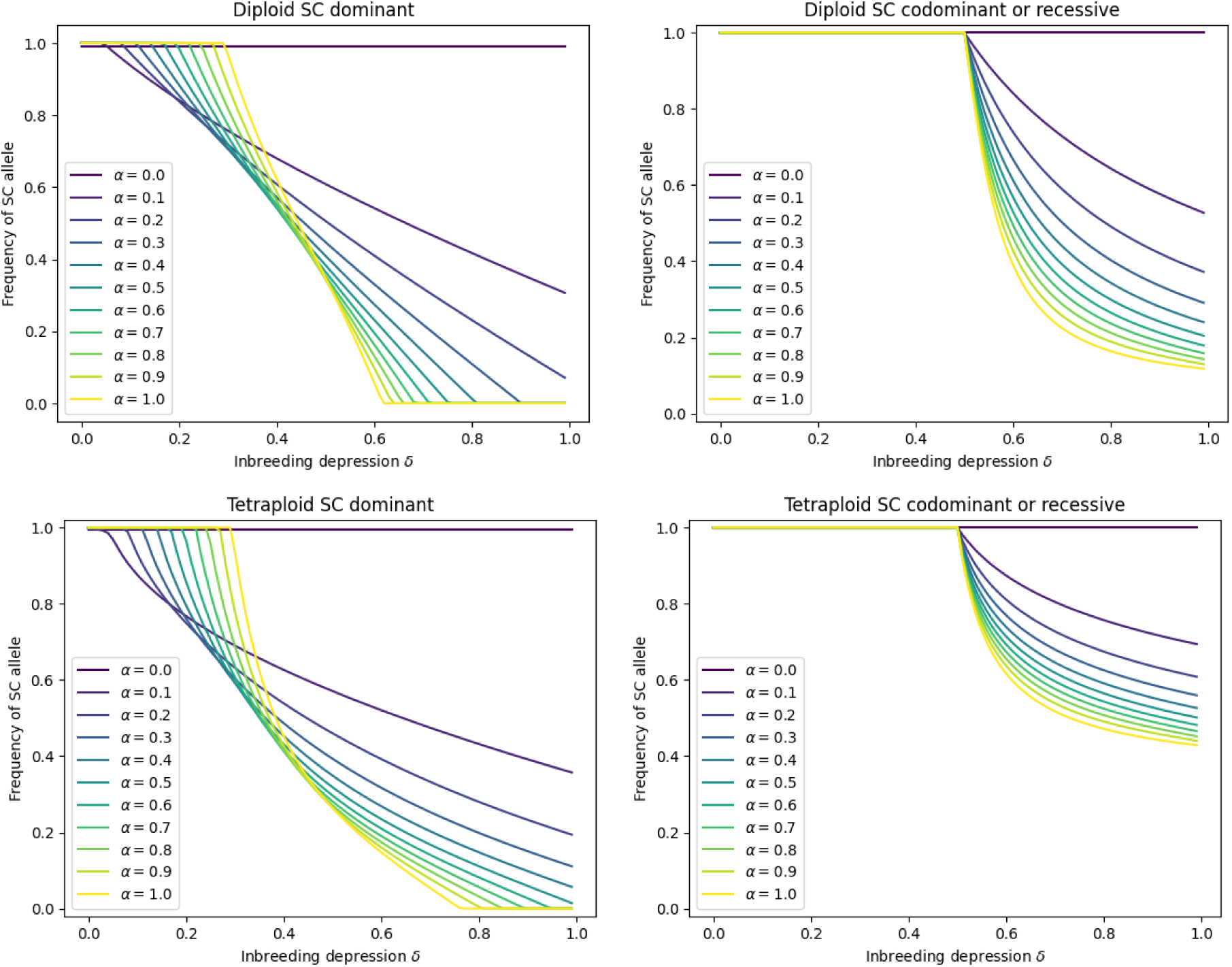
SI codominant. Frequency of SC allele at equilibrium as a function of inbreeding depression for different values of the self-pollen rate (represented by different colored lines). Results obtained by recursions on the analytical models for diploids when *n* = 20 (top) and for tetraploids when *n* = 40 (bottom), when the SC allele is dominant (left) or codominant/recessive (right).

**Figure 4:**
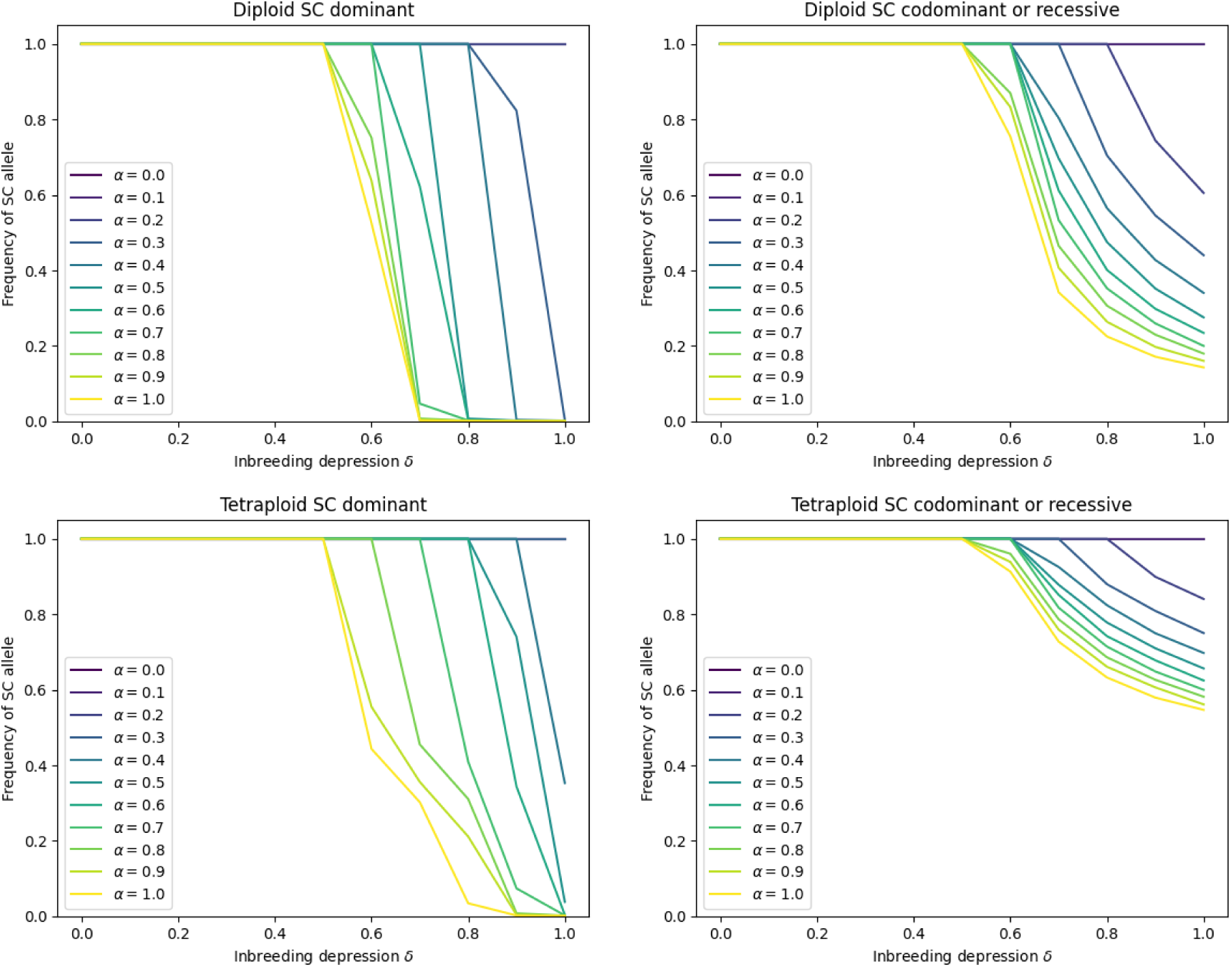
SI codominant. Frequency of SC allele at equilibrium as a function of inbreeding depression for different values of the self-pollen rate (represented by different colored lines) when *N*_pop_ = 1000. Results obtained using the individual-based model (10 runs) when inbreeding depression *δ* is fixed for diploids (top) and tetraploids (bottom) when the SC allele is dominant (left) or codominant/recessive (right).

Although simulation results are consistent with analytical predictions for the minimum inbreeding depression preventing SC maintenance in the population, the two approaches differ greatly regarding SC fixation versus coexistence with SI alleles (represented by the orange curves in Figure 2). Coexistence predominates in the analytical model, whereas simulations more frequently lead to SC fixation, especially when the SC allele is dominant over all SI alleles (Figures 3 and 4). As these differences concern both diploids and tetraploids, they are thoroughly described later in this section.

#### In tetraploids

When the SC allele is dominant over all SI alleles, Figure 2 shows that SSI is fully maintained only over a limited range of parameters, while fixation or coexistence between SC and SI alleles occurs more broadly (Figure 3). Simulations results are consistent with analytical predictions when inbreeding depression is fixed in the individual-based model (Figures 2, 3 and 4), although simulations tend to show more invasion than coexistence. In contrast, when inbreeding depression is controlled by the mutation rate *U* , Figure S6 exhibits a reversal of the expected dynamics, with the SC allele reaching fixation for high inbreeding depression levels. This likely reflects the high deleterious mutation rates required in tetraploids to achieve such levels (e.g., *U* = 10 for *δ* ≈ 1), due to the more effective purging of deleterious alleles in polyploids. These high mutation rates reduce overall fitness to values that are numerically indistinguishable from zero and strongly compromise the effectiveness of the selection process. However, because the model assumes a constant population size, populations do not undergo extinction under these conditions, leading to biologically unrealistic dynamics. As fitness becomes extremely low, compatible mates with sufficiently high fitness become scarce, and the number of S-alleles declines, further limiting mating opportunities and favoring self-compatible individuals, which do not depend on mate availability for reproduction.

Both approaches suggest that the SC mutant is maintained in a tetraploid population when the SC allele is codominant with or recessive to all SI alleles regardless of the value of inbreeding depression and for all selfing rates (see Figures 2, 3 and 4, and Section A.2.2 of the appendix). In addition, Figures 3 and 4 reveal that the SC allele is fixed for a wide range of parameters, and is always present at high frequencies in the population at equilibrium for all parameter values.

#### Diploids VS Tetraploids

These results highlight both differences and similarities in the conditions under which a mutant SC allele can be maintained in diploids versus tetraploids. Figure 2 highlights that, for a given population size (*N*_pop_ = 1000), the SSI system is maintained in diploids under lower values of the self-pollen rate *α* and lower levels of inbreeding depression than in tetraploids, when the SC allele is dominant. More generally, for the same set of parameters, Figures 3 and 4 indicate that the frequency of the SC allele at equilibrium is higher in tetraploid than in diploid populations, in both dominance scenarios. Overall, these results suggest that SSI systems are more difficult to maintain in tetraploids than in diploids due to rarer selfing events and a stronger advantage in outcrossing.

Figures 3 and 4 show that, in both diploids and tetraploids, inbreeding depression and the rate of self-pollen have an impact on the frequency of the SC mutant allele in the population at equilibrium. Figures 3 and 4 reveal that the equilibrium frequency of the SC allele decreases as inbreeding depression increases. Moreover, under high inbreeding depression, increasing the self-pollen rate further reduces the frequency of the SC allele at equilibrium. In contrast, under low inbreeding depression, the equilibrium frequency of the SC allele increases with the self-pollen rate, as selfing is only weakly constrained by inbreeding depression, which favors the spread of the SC allele. This is observed in both tetraploid and diploid populations, regardless of the dominance of the SC allele. Additionally, for diploids and tetraploids, the impact of inbreeding depression and of the self-pollen rate is stronger when the SC allele is dominant than when it is codominant or recessive to all SI alleles, due to higher frequencies of self-compatible individuals in the population. Moreover, Figures S4 and S5 indicate that the SSI system is maintained for a wider range of parameters when the number of S-alleles in the population (directly related to the population size) increases, which is expected for both ploidy levels.

In both diploids and tetraploids, coexistence is mainly observed in the analytical results, while simulation results exhibit fixation of the SC allele in the population for a wider range of parameters (orange curves in Figure 2). These discrepancies can be attributed to key differences between the two models. In the analytical model, the equilibrium number of SI alleles *n* is fixed, while in the individual-based simulations *n* declines as the SC allele spreads, leading to an increase in the frequency of the SC allele. Under low inbreeding depression, the frequency of the SC allele is ultimately pushed to levels high enough for the remaining SI alleles to be lost through genetic drift as fitness differences are weak. However, at higher levels of inbreeding depression, the loss of residual SI alleles by drift becomes increasingly unlikely. Individuals carrying an SI allele are mostly, or even exclusively (in the codominant/recessive case), produced by outcrossing, which confers a substantial fitness advantage when inbreeding depression is strong, thus maintaining SI alleles in the population. In addition, S-alleles in the individual-based simulations undergo recurrent mutations from SI to SC, which may promote the fixation of the SC allele when simulations are run over a large number of generations.

### 3.2 SI alleles structured in dominance classes

We consider the scenario where SI alleles are distributed into four possible dominance classes and investigate, as in the codominant case, the impact of the dominance of the SC allele, as well as the impact of the ploidy level, on the maintenance or breakdown of the SSI system. The SC allele either belongs to one of the SI dominance classes, or is dominant over all SI alleles (class 5). This analysis is mostly based on the results obtained using individual-based simulations in which the level of inbreeding depression *δ* is fixed, rather than controlled by deleterious mutations, to avoid any numerical issues. Analytical predictions are also used for comparison in the diploid case.

#### In diploids

Figures 5 and 6 show that, under high inbreeding depression (above 0.7), there exist parameter sets in which the SSI system is maintained across all dominance scenarios for the SC allele. In contrast, when SI alleles are codominant, the SC allele is always maintained in the population when it is codominant or recessive (Figure 2), suggesting that structuring SI alleles into dominance classes favors the maintenance of the SSI system. When *S_C_* belongs to class 5, the same pattern as in the codominant SI case with a dominant SC allele is recovered. Additionally, Figure 5 indicates that SSI is more readily maintained when the SC allele belongs to a higher dominance class in the deterministic model, except for the lowest dominance classes (classes 1–2) in simulations. However, Figure 6 reveals no clear pattern in diploids, suggesting that the dominance class of the SC allele has a limited effect on its equilibrium frequency. While SSI appears to be maintained at lower levels of inbreeding depression when the SC allele belongs to a higher dominance class in Figure 5, Figure 6 shows that, for some parameter sets, the equilibrium frequency of the SC allele is higher in these classes, leading to conflicting trends (e.g., when *α* = 0.1 or 0.2 and *δ* = 1). Overall, these results do not provide a clear conclusion regarding the impact of the SC allele’s dominance class on the maintenance or breakdown of SSI in diploid populations.

**Figure 5:**
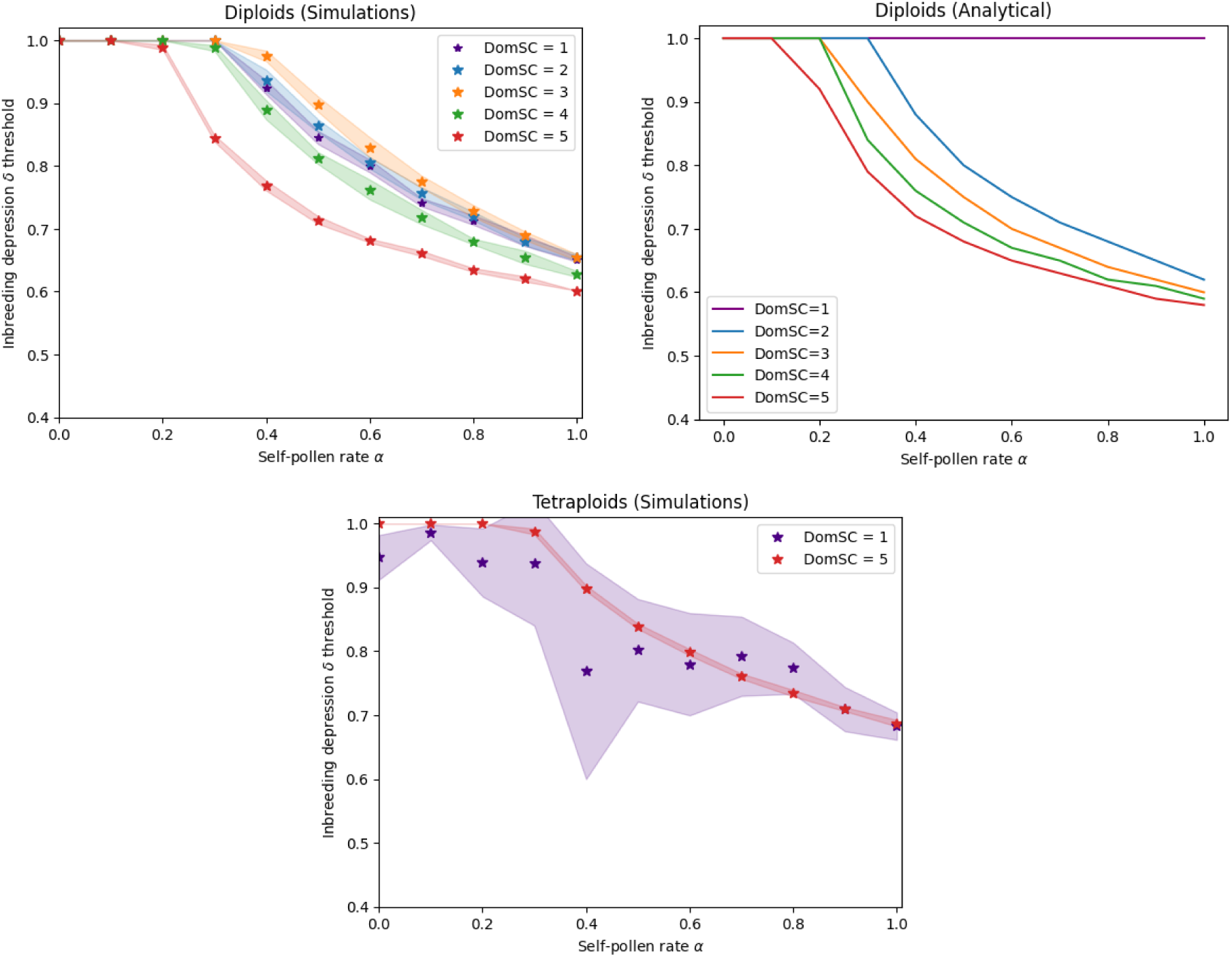
SI alleles in dominance classes. Minimal inbreeding depression necessary to prevent the maintenance of an SC mutant allele into an initially SI population with fixed inbreeding depression *δ*. Results obtained using individual-based simulations (10 runs) represented by colored stars with 95% confidence intervals, and analytical predictions for diploids (A17) represented by colored lines. The number of SI alleles in the deterministic model (*n* = 15) and their distribution into dominance classes ([1, 2, 4, 8]) were chosen by computing the average values observed in simulations before introducing the SC allele. Comparison between diploids (top) and tetraploids (bottom) under different dominance scenarios, where the SC allele either belongs to one of the dominance classes, or is dominant over all SI alleles (class 5).

**Figure 6:**
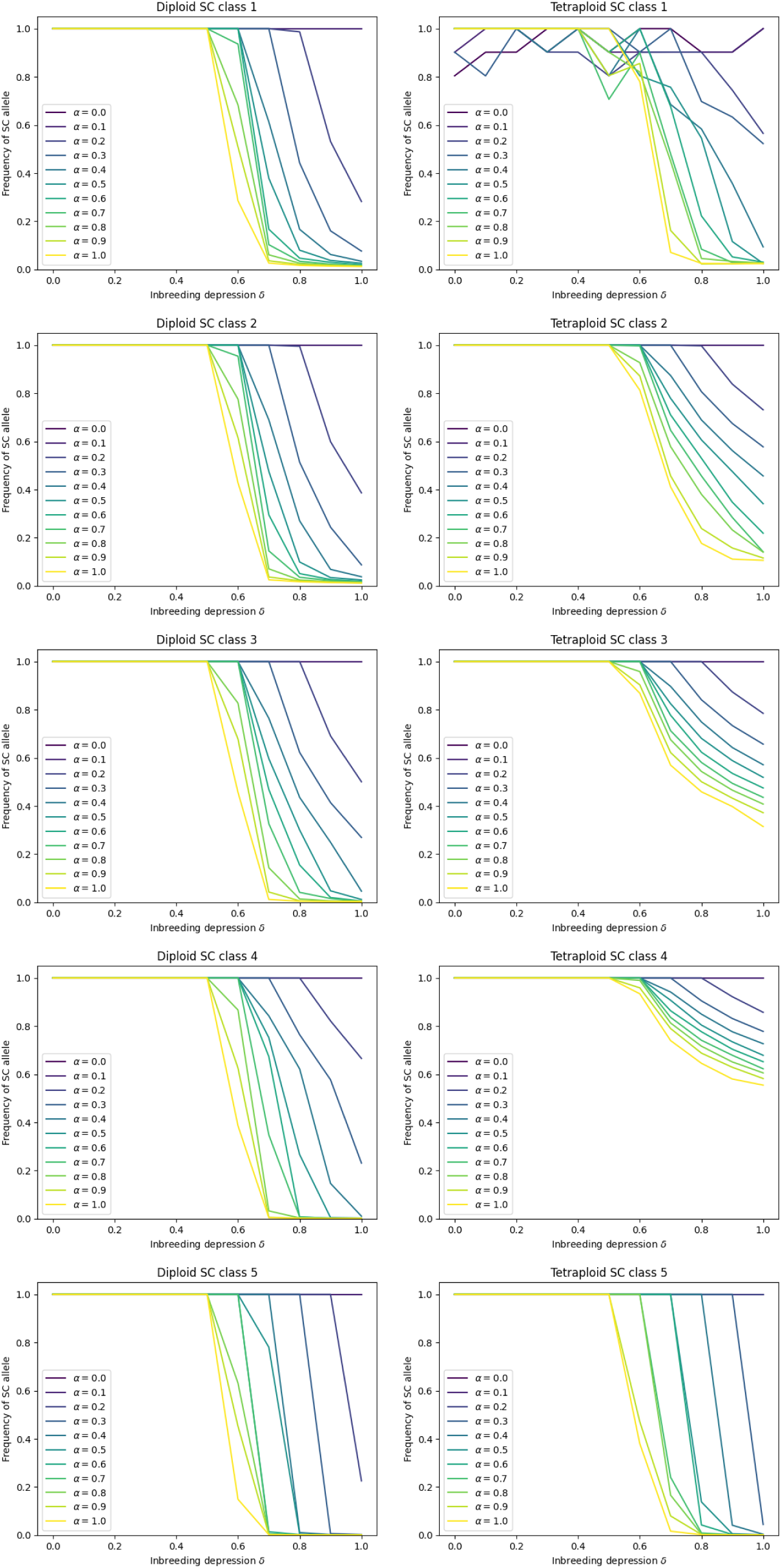
SI alleles in dominance classes. Frequency of SC allele at equilibrium as a function of inbreeding depression for different values of the self-pollen rate (represented by different colored lines) when *N*_pop_ = 1000. Results obtained using the individual-based model (10 runs) with fixed inbreeding depression *δ*, for diploids (left) and tetraploids (right) when the SC allele belongs to different classes of dominance.

Simulations and analytical predictions show similar patterns for highly dominant SC alleles, although some discrepancies remain (Figure 5). Analytical results predict lower inbreeding depression thresholds for higher dominance classes, with a monotonic decrease as the SC allele dominance increases. In contrast, simulations produce intermediate thresholds for the lowest dominance classes, between those of classes 3 and 4. These discrepancies could be attributed to differences between the models. Because individual-based simulations are stochastic, we need to define a threshold to determine whether the SSI system is maintained (when the frequency of the SC allele remains below 0.05). However, the SC allele may persist at a very low frequency without being completely eliminated from the population at equilibrium. Moreover, the deterministic model requires fixed values for both the number of S-alleles and their distribution across dominance classes, whereas these quantities fluctuate in individual-based simulations. Notably, even though up to four dominance classes are possible for SI alleles, only the three most dominant are maintained at equilibrium in some simulations, the most recessive class being lost once the number of SI alleles stabilizes. The observed discrepancies are likely due to the difficulty of reproducing simulation conditions within the deterministic framework.

#### In tetraploids

A clear pattern emerges when the SC allele belongs to one of the SI dominance classes: its frequency at equilibrium increases with its dominance class (Figures 5 and 6). In particular, Figure 6 shows that the SC allele is consistently maintained in the population when it belongs to classes 2,3 and 4. In contrast, when the SC allele is dominant over all SI alleles (class 5), it is absent from the population at higher levels of inbreeding depression. Results for class 1 (Figures 5 and 6) exhibit strong stochasticity, probably because SI alleles are not consistently maintained in this dominance class across simulations runs.

#### Diploids versus Tetraploids

Figures 5 and 6 highlight clear differences in the maintenance or breakdown of SSI systems between diploid and tetraploid populations. First, Figure 5 shows that, while SSI can be maintained across all dominance scenarios for the SC allele in diploids under certain parameter sets, this is only observed in tetraploids when the SC allele belongs to either class 1 or class 5. In other cases, the SC allele is always maintained in tetraploids (Figure 6). Moreover, even when SSI is fully maintained in tetraploids, it occurs over a smaller range of parameter values than in diploids for a given dominance scenario, indicating that the SC allele is more likely to be maintained in tetraploid than in diploid populations, leading to the loss of SSI.

Another striking difference, visible in Figure 6, is that diploids exhibit a weaker dependence on the dominance class of the SC allele than tetraploids, where the differences between the dominance scenarios are more significant. This suggests that the dominance class in which the SC allele emerges has a stronger impact on its maintenance in tetraploid populations. Except when the SC allele is dominant over all SI alleles, diploids and tetraploids show distinct dynamics with respect to the dominance class of the SC allele. In tetraploids, the likelihood of invasion increases with the dominance class of the SC allele, a pattern that is not observed in diploids.

## 4 Discussion

### SSI breakdown in tetraploids versus diploids

Our results suggest that the break-down of SSI systems is more likely in tetraploids than in diploids. For a given parameter set, the SC allele consistently reaches higher equilibrium frequencies in tetraploids across all scenarios, while its complete loss occurs only rarely. This may appear counterintuitive because tetraploid populations maintain a larger number of S-alleles (here, *n* = 30–40 on average) than diploids (*n* = 20) for the same population size, which should in principle hinder the invasion of a mutant SC allele. However, tetraploid individuals carry four S-alleles, making the SSI system more restrictive and reducing the number of compatible mates, particularly when SI alleles are codominant. Under these conditions, individuals carrying one or more SC alleles gain a substantial advantage in outcrossing because they express fewer SI specificities and therefore have access to more potential mates.

In our models, ploidy levels were assumed to remain constant over time. Consequently, we did not consider scenarios in which tetraploids could appear within initially diploid populations, for example through the production of unreduced gametes as investigated in Douet et al. (2026). Extending the model to allow changes in ploidy would make it possible to examine how dominance relationships and inbreeding depression influence tetraploid establishment. Although SSI appears less stable in tetraploids than in diploids, our results suggest that the SC allele often spreads through an outcrossing advantage rather than through increased selfing, raising questions about the causal relationship between polyploidy and the evolution of self-fertilization. This is consistent with recent evidence that autopolyploidy can exacerbate dominance masking under negative frequency-dependent selection and alter sporophytic self-incompatibility dynamics in *Arabidopsis* (Vekemans et al., 2026), although the authors did not find evidence for a breakdown of SSI in autotetraploid populations.

### Inbreeding depression and deleterious mutations

Across all scenarios, the minimum level of inbreeding depression required to maintain SSI was lower in diploids than in tetraploids. Inbreeding depression therefore strongly contributes to the maintenance of self-incompatibility at both ploidy levels, although its effects vary according to dominance relationships and appear to be more pronounced in diploids. These findings are consistent with previous theoretical studies showing that high levels of inbreeding depression favor the maintenance of self-incompatibility when deleterious mutations are mildly harmful, whereas the purging of highly deleterious or lethal mutations may facilitate the spread of SC alleles (Porcher and Lande, 2005b; Gervais et al., 2014). Because lethal mutations were not included in our model, our results specifically describe the effects of mildly deleterious mutations on the maintenance and breakdown of SSI while highlighting the additional roles of ploidy and dominance.

When inbreeding depression was determined by the deleterious mutation rate, some tetraploid simulations produced numerical issues, with particular parameter combinations leading to zero fitness values. These outcomes resulted from the very high mutation rates required to generate strong inbreeding depression. Such mutations rates may be unrealistic in natural tetraploid populations, especially because empirical studies generally report lower levels of inbreeding depression (Clo and Kolář, 2022). Moreover, excessively high levels of inbreeding depression may itself increase the risk of extinction (Abu Awad and Billiard, 2017). These limitations suggest caution when interpreting results obtained under strong mutational loads.

### Dominance of the SC allele

Our results show that dominance relationships strongly influence the maintenance of SSI. Across all scenarios, SSI is more easily maintained when the SC allele is dominant over all SI alleles. A likely explanation is that dominant SC alleles are more exposed to inbreeding depression as they increase the frequency of selfing events, whereas more recessive SC alleles restrict selfing and therefore experience weaker fitness costs associated with inbreeding.

This pattern is particularly clear when all SI alleles are codominant. In that case, all individuals carrying at least one SC allele are self-compatible when *S_C_* is dominant over all SI alleles. Note that the case in which *S_C_* is dominant is theoretical, as codominance among all SI alleles suggests the absence of a dominance modifier at the SI locus (Durand et al., 2014), though it still provides valuable insight into the mechanisms underlying the breakdown of SSI. In contrast, when *S_C_* is codominant or recessive, only individuals homozygous for *S_C_* are self-compatible, greatly reducing opportunities for self-fertilization. Under these conditions, the spread of the SC allele occurs mainly through outcrossing, where it confers an advantage by reducing the number of expressed SI specificities, thus increasing access to compatible mates. In tetraploids, this effect is even stronger because self-fertilization is restricted to individuals carrying four copies of *S_C_*, further reducing the impact of inbreeding depression.

### Partial dominance of the SC allele

When SI alleles are organized into dominance classes, a different pattern emerges. If the SC allele is codominant with at least some SI alleles, increasing the dominance of *S_C_* facilitates SSI breakdown, particularly in tetraploids. In this situation, individuals carrying the SC allele gain a stronger outcrossing advantage because they express fewer SI specificities in pollen, while self-fertilization remains constrained by codominance with several SI alleles. Consequently, SC alleles benefit simultaneously from improved mate availability and limited exposure to inbreeding depression.

These results are consistent with previous theoretical work showing that SC alleles may possess a substantial invasion advantage (Charlesworth, 1988). However, our results further suggest that this advantage is maximized when the SC allele is dominant over only a subset of SI alleles rather than over all of them. This interpretation is also compatible with empirical observations showing that highly dominant SC alleles can lead to an immediate transition towards selfing following hybridization or polyploidization (Novikova et al., 2023; Duan et al., 2024). By contrast, diploid populations show less consistent responses to dominance-class structure, suggesting that dominance relationships have a stronger effect on SSI breakdown in tetraploids than in diploids. Overall, our results indicate that SC invasion often relies more on enhanced outcrossing success than on increased self-fertilization.

### Coexistence or invasion of the SC allele

Analytical predictions revealed coexistence between SC and SI alleles more frequently than individual-based simulations, especially for lower levels of inbreeding depression. These discrepancies likely arise from the assumptions underlying analytical models, which neglect demographic stochasticity and fluctuations in S-allele diversity. By contrast, simulations explicitly incorporate these processes and therefore allow stochastic loss or fixation of alleles.

Whether the SC allele invades the population or coexists with SI alleles is determined by several factors that either favor the spread of the SC allele or prevent it. Selfing provides an automatic transmission advantage and reproductive assurance, especially in environment where compatible mates are rare, for example due to low S-alleles diversity or pollen limitation, which favors SC invasion unless inbreeding depression is sufficiently strong (Porcher and Lande, 2005b; Vallejo-Marín and Uyenoyama, 2004; Busch and Schoen, 2008). However, even though inbreeding depression can prevent the invasion of the SC allele, this barrier may become weaker if deleterious mutations are purged over time (Gervais et al., 2014). Together, these effects can lead to either stable coexistence between SC and SI phenotypes or complete replacement of SI alleles by the SC allele, which have both been observed in natural populations (De Cauwer et al., 2021; Igic et al., 2008).

### S-allele diversity

Beyond the effects of deleterious mutations, several factors may influence the diversity of S-alleles maintained within populations, including population size and pollen limitation. Previous work showed that larger populations should maintain a greater number of S-alleles because negative frequency-dependent selection is more effective and genetic drift is weaker, thus increasing overall S-locus diversity (Schierup et al., 1997). Therefore, increasing the population size makes the breakdown of SSI systems more difficult, as shown in our results. Likewise, pollen limitation can alter the selective balance between SI and SC individuals by increasing the reproductive costs associated with self-incompatibility (Vallejo-Marín and Uyenoyama, 2004; Busch and Schoen, 2008).

The structure of dominance interactions can also affect S-allele diversity. In simulations with dominance classes, the most recessive class was frequently lost. A similar pattern was reported by Schierup et al. (1997), who showed that under codominance in the pistil and dominance in the pollen, the most recessive alleles may be lost despite negative frequency-dependent selection and may also have lower invasion probabilities. However, this tendency can be reversed under pollen-limited conditions, where stronger selection favors the persistence of even the most recessive alleles (Vekemans et al., 1998). These results suggest that the maintenance of S-allele diversity depends not only on balancing selection, but also on ecological and demographic conditions.

### Model assumptions on dominance

The patterns reported here may vary under alternative dominance architectures. We assumed that dominance relationships are expressed only in pollen, whereas S-alleles in the pistil were considered strictly codominant. This assumption is a reasonable approximation based on the observation within *Brassicaceae* of a predominance of dominance relationships in the pollen and of codominance in the pistil (Stevens and Kay, 1989; Hatakeyama et al., 1998; Llaurens et al., 2008), but it does not capture the full diversity of SSI systems. Dominance relationships in the pistil (Kowyama et al., 1994; Brennan et al., 2006), incomplete codominance among SI alleles (Hatakeyama et al., 1998), or variation in the number of dominance classes (Fujii and Takayama, 2018) have all been described in natural systems and could lead to different outcomes between SC invasion and SSI maintenance.

We also assumed the introduction of a single SC allele. In natural populations, multiple SC alleles may arise independently, potentially inheriting dominance relationships from ancestral S-haplotypes that subsequently became non-functional (Tsuchimatsu et al., 2010). Although each SC allele might individually follow the dynamics predicted here, interactions among several SC alleles could generate outcomes that differ from those observed in our model. Exploring these more complex genetic architectures represents an important direction for future theoretical studies.

## 5 Conclusion

Overall, our results show that SSI systems are harder to maintain in tetraploid than in diploid populations, even though tetraploids maintain a higher diversity of S-alleles. This is mainly because carrying four S-alleles makes the SSI system more restrictive, which gives the SC mutant allele a stronger outcrossing advantage. Moreover, inbreeding depression remains an important barrier to the spread of an SC allele, but our results show that the evolution of self-compatibility depends on more than just the balance between selfing and inbreeding depression. It is also influenced by ploidy level and by the dominance relationships at the S-locus. In particular, the dominance of the SC allele plays a key role in determining whether SSI is maintained, whether the SC allele coexists with SI alleles, or whether it invades the population. Dominance relationships among SI alleles also matter, and different outcomes can be expected depending on the structure of SI dominance classes. Taken together, these results highlight the role of polyploidy and dominance in the evolution of mating strategies.

## Data availability

The simulation models and the *Wolfram Mathematica* notebooks are available at https://gitlab.univ-lille.fr/eep_lab/diane_douet/SSI_breakdown.

## Acknowledgments

We thank Justin Dallant for helpful advice on code optimization in *Julia*, and Mathieu Genete for his assistance in making scripts and notebooks available on GitLab.

