## Supplementary material for "Breakdown of sporophytic self-incompatibility: Diploids versus tetraploids"

### A Supplementary Material

#### A.1 Analytical models

This section provides the recursion equations of the analytical models in the case where all SI alleles are codominant for diploid (in Section A.1.1) and tetraploid populations (in Section A.1.2), presented in Sections 2.1.1 and 2.1.2 of the main text respectively. Section A.1.3 also gives the recursion equations of the deterministic model for diploids in the case where SI alleles are distributed into four dominance classes, presented in Section 2.2.1 of the main text.

##### A.1.1 Diploids: SI codominant

We denote by $x_1$ and $x_2$ the frequencies of individuals with genotype $S_C S_C$ produced by selfing and outcrossing, respectively. Similarly, we denote by $x_3$ and $x_4$ the frequencies of $S_i S_C$ individuals produced by selfing and outcrossing, respectively. We also denote

$$x_0 = 1 - x_1 - x_2 - x_3 - x_4 \tag{A1}$$

the frequency of $S_i S_j$ individuals. We assume that selfed individuals experience inbreeding depression, such that $W_0 = W_2 = W_4 = 1$, while $W_1 = W_3 = 1 - \delta$. The mean fecundity is therefore expressed by $\overline{W} = 1 - \delta(x_1 + x_3)$. The selfing rates for each group are denoted by $a_1$, $a_2$, $a_3$ and $a_4$. We use the symbol $\neg$ to denote logical negation ("not").

**$S_C$ codominant or recessive.** In this paragraph, we assume that the $S_C$ allele is codominant or recessive in the pollen with respect to all SI alleles. Note that in that case, only $S_C S_C$ individuals are self-compatible.

Therefore, assuming that all SI alleles are present at equal frequency, we define the following frequencies of gametes in the pollen pool, with $n$ the number of SI alleles maintained in the population before introducing the SC allele:

- The frequency of gametes produced by $S_C S_C$ individuals: $q_{CC} = \frac{1}{\overline{W}}(W_1 x_1 + W_2 x_2)$;

- The frequency of gametes with genotype $S_C$: $q = q_{CC} + \frac{1}{2\overline{W}}(W_3 x_3 + W_4 x_4)$;

- Given $i$, the frequency of gametes produced by $S_C S_k$ individuals for all $k \neq i$: $q_{C,\neg i} = \frac{1}{\overline{W}} \frac{(n-1)}{n}(W_3 x_3 + W_4 x_4)$;
- Given $i$ and $j$, the frequency of gametes produced by $S_C S_k$ individuals, for all $k \neq i, j$: $q_{C,\neg ij} = \frac{1}{\overline{W}} \frac{n-2}{n}(W_3 x_3 + W_4 x_4)$;
- Given $i$, the frequency of gametes produced by $S_k S_l$ individuals for all $k, l \neq i$: $q_{\neg i,\neg i} = \frac{x_0}{\overline{W}} \frac{n-2}{n}$;
- Given $i$ and $j$, the frequency of gametes produced by $S_k S_l$ individuals for all $k, l \neq i, j$: $q_{\neg ij,\neg ij} = \frac{x_0}{\overline{W}} \frac{(n-2)(n-3)}{n(n-1)}$.

All expressions are obtained using the same combinatorial construction, by decomposing the event into the frequency of the relevant pollen genotype among the pollen pool multiplied by the conditional probability that it satisfies the required allelic condition. To illustrate how these frequencies are derived, consider $q_{C,\neg i}$ as an example. The frequency of pollen carrying genotype $S_C S_k$ (for any $k$) is $\frac{1}{\overline{W}}(W_3 x_3 + W_4 x_4)$, which is multiplied by the conditional probability that such pollen does not carry allele $S_i$, namely $\frac{n-1}{n}$.

The simplified frequencies at the next generation are given by the following equations:

$$\begin{aligned}
\overline{W}x_1' =& W_1 a_1 x_1 + W_2 a_2 x_2, \\
\overline{W}x_2' =& W_1(1-a_1)x_1 q + W_2(1-a_2)x_2 q \\
&+ \frac{1}{2}[W_3(1-a_3)x_3 + W_4(1-a_4)x_4] \frac{q_{CC} + \frac{1}{2}q_{C,\neg i}}{q_{CC} + q_{C,\neg i} + q_{\neg i,\neg i}}, \\
\overline{W}x_3' =& 0, \\
\overline{W}x_4' =& W_1(1-a_1)x_1(1-q) + W_2(1-a_2)x_2(1-q) \\
&+ \frac{1}{2}W_3(1-a_3)x_3 + \frac{1}{2}W_4(1-a_4)x_4 + x_0 \frac{q_{CC} + \frac{1}{2}q_{C,\neg ij}}{q_{CC} + q_{C,\neg ij} + q_{\neg ij,\neg ij}}.
\end{aligned} \tag{A2}$$

Each recursion equation is obtained by summing the contributions of all reproductive events producing offspring in the corresponding group. As an illustration, consider the recursion for $x_2'$, corresponding to outcrossed $S_C S_C$ individuals. This group receives contributions from:

- outcrossed $S_C S_C$ mothers fertilized by pollen with genotype $S_C$;

- outcrossed $S_C S_i$ mothers, which transmit the SC allele with probability 1/2 and are fertilized by compatible pollen carrying the SC allele. The denominator represents the total frequency of compatible pollen and therefore accounts for competition among all compatible pollen grains.

$S_C$ **dominant.** In this paragraph, we assume that the SC allele is dominant over all SI alleles in the pollen. In that case, both $S_C S_C$ and $S_C S_i$ individuals are self-compatible for all $i$. As $S_i S_i$ individuals appear in the population, we define new expressions for the following frequencies:

- The frequency of gametes produced by $S_C S_k$ individuals for all $k$:
  $q_{CI} = \frac{1}{\overline{W}}(W_3 x_3 + W_4 x_4)$;
- Given $i$, the frequency of gametes produced by $S_k S_l$ individuals for all $k, l \neq i$:
  $p_{\neg i, \neg i} = \frac{x_0}{\overline{W}} \frac{(n-1)^2}{n^2}$;
- Given $i$ and $j$, the frequency of gametes produced by $S_k S_l$ individuals for all $k, l \neq i, j$:
  $p_{\neg ij, \neg ij} = \frac{x_0}{\overline{W}} \frac{(n-2)^2}{n^2}$.

Note that $S_i S_i$ individuals can appear through selfing and therefore suffer from inbreeding depression. Inbreeding depression in these individuals is neglected as the exact and approximate models lead to very similar outcomes (see *Wolfram Mathematica* Notebook for more details and Figure S1). The frequencies $q$ and $q_{CC}$ defined above are unchanged and are therefore used in this model.

The frequencies in the next generation are given by the following equations:

$$\begin{aligned}
\overline{W} x_1' =& W_1 a_1 x_1 + W_2 a_2 x_2 + \frac{1}{4} W_3 a_3 x_3 + \frac{1}{4} W_4 a_4 x_4, \\
\overline{W} x_2' =& W_1 (1 - a_1) x_1 q + W_2 (1 - a_2) x_2 q \\
&+ \frac{1}{2} [W_3 (1 - a_3) x_3 + W_4 (1 - a_4) x_4] \frac{q}{q_{CC} + q_{CI} + p_{\neg i, \neg i}}, \\
\overline{W} x_3' =& \frac{1}{2} W_3 a_3 x_3 + \frac{1}{2} W_4 a_4 x_4, \\
\overline{W} x_4' =& W_1 (1 - a_1) x_1 (1 - q) + W_2 (1 - a_2) x_2 (1 - q) + \frac{1}{2} W_3 (1 - a_3) x_3, \\
&+ \frac{1}{2} W_4 (1 - a_4) x_4 + x_0 \frac{q}{q_{CC} + q_{CI} + p_{\neg ij, \neg ij}}.
\end{aligned} \tag{A3}$$

##### A.1.2 Tetraploids: SI codominant

We denote by $x_{1s}$, $x_{2s}$, $x_{3s}$ and $x_{4s}$ the frequencies of individuals with 1,2,3 or 4 SC alleles produced by selfing. Similarly, we denote by $x_{1c}$, $x_{2c}$, $x_{3c}$ and $x_{4c}$ the frequencies of individuals with 1,2,3 or 4 SC alleles produced by outcrossing. We also denote:

$$x_0 = 1 - x_{1s} - x_{2s} - x_{3s} - x_{4s} - x_{1c} - x_{2c} - x_{3c} - x_{4c} \tag{A4}$$

the frequency of $S_iS_jS_kS_l$ individuals (with no SC alleles). We assume that selfed individuals experience inbreeding depression, such that $W_0 = W_{1c} = W_{2c} = W_{3c} = W_{4c} = 1$, while $W_{1s} = W_{2s} = W_{3s} = W_{4s} = 1 - \delta$. The mean fecundity is therefore expressed by $\overline{W} = 1 - \delta(x_{1s} + x_{2s} + x_{3s} + x_{4s})$. The selfing rates for self-compatible groups are denoted by $a_{1s}$, $a_{2s}$, $a_{3s}$ and $a_{4s}$ for selfed individuals, and $a_{1c}$, $a_{2c}$, $a_{3c}$ and $a_{4c}$ for outcrossed individuals.

**$S_C$ codominant or recessive.** In this paragraph, we consider that the SC allele is codominant with or recessive to all SI alleles. In that case, only fully homozygous individuals at the S-locus ($S_CS_CS_CS_C$ individuals) are self-compatible. For example, a pollen with genotype $S_CS_C$ that come from an $S_CS_CS_iS_j$ individual will not be able to fertilize any mother that carries alleles $S_i$ or $S_j$.

For $k \in [\![0, 3]\!]$ and $l \in [\![0, 4]\!]$, the frequency of gametes produced by an individual with $k$ SC alleles, in which all the SI alleles are different from $l$ given S-alleles, is given by:

$$f_{kl} = \frac{1}{\overline{W}}(W_{kc}x_{kc} + W_{ks}x_{ks})\prod_{i=0}^{3-k}\frac{n-l-i}{n-i}. \tag{A5}$$

In addition, we define $f_4$ the frequency of gametes produced by individuals with four SC alleles, that are compatible with all individuals in the population, as:

$$f_4 = \frac{1}{\overline{W}}(W_{4c}x_{4c} + W_{4s}x_{4s}). \tag{A6}$$

For example, the frequency of gametes produced by an $S_CS_CS_iS_j$ individual (with two

SC alleles), in which $i, j \neq I, J, K$ for given $I, J, K$ and for all $i, j$ is given by:

$$f_{23} = \frac{1}{\overline{W}}(W_{2c}x_{2c} + W_{2s}x_{2s})\frac{(n-3)(n-4)}{n(n-1)}. \tag{A7}$$

The last factor corresponds to the probability that the two distinct SI alleles (as $S_iS_i$ individuals cannot appear in this case) are both different from $I$, $J$, and $K$. Among the $n(n-1)$ possible pairs of distinct SI alleles, the first allele can be chosen in $n-3$ ways (excluding $I$, $J$, and $K$), and the second in $n-4$ ways among the remaining admissible alleles, leading to $(n-3)(n-4)$ favorable pairs.

With the above frequencies, we define the frequency of pollen with genotypes $S_CS_C$, $S_CS_i$ and $S_iS_j$ for all $i, j$ in the population, respectively, that are compatible with a maternal parent that has $k$ SI alleles:

$$\begin{aligned}
p_{CC,k} &= f_4 + \frac{1}{2}f_{3k} + \frac{1}{6}f_{2k}, \\
p_{CI,k} &= \frac{1}{2}f_{3k} + \frac{2}{3}f_{2k} + \frac{1}{2}f_{1k}, \\
p_{II,k} &= \frac{1}{6}f_{2k} + \frac{1}{2}f_{1k} + f_{0k}.
\end{aligned} \tag{A8}$$

For example, consider $p_{CC,2}$, the frequency of compatible $S_CS_C$ pollen for a maternal parent carrying two SI alleles (e.g., genotype $S_CS_CS_iS_j$):

$$p_{CC,2} = f_4 + \frac{1}{2}f_{32} + \frac{1}{6}f_{22}. \tag{A9}$$

Such pollen can originate from three types of paternal individuals:

- Individuals with four SC alleles, with frequency $f_4$, which always produce compatible $S_CS_C$ pollen.
- Individuals with three SC alleles whose SI allele is different from $S_i$ and $S_j$, with frequency $f_{32}$. These individuals produce $S_CS_C$ pollen with probability $\frac{1}{2}$.
- Individuals with two SC alleles whose two SI alleles are both different from $S_i$ and $S_j$, with frequency $f_{22}$. These individuals produce $S_CS_C$ pollen with probability $\frac{1}{6}$.

We also define the frequency of pollen compatible with a maternal parent that has $k$ SI alleles as follows:

$$D_k = f_4 + f_{3k} + f_{2k} + f_{1k} + f_{0k}. \tag{A10}$$

The frequencies in the next generation are given by the following equations:

$$\begin{aligned}
\overline{W}x'_{4s} =& W_{4c}x_{4c}a_{4c} + W_{4s}x_{4s}a_{4s}, \\
\overline{W}x'_{3s} =& 0, \\
\overline{W}x'_{2s} =& 0, \\
\overline{W}x'_{1s} =& 0, \\
\overline{W}x'_{4c} =& [W_{4c}x_{4c}(1-a_{4c}) + W_{4s}x_{4s}(1-a_{4s})]p_{CC,0} \\
&+ \frac{1}{2}[W_{3c}x_{3c}(1-a_{3c}) + W_{3s}x_{3s}(1-a_{3s})]\frac{p_{CC,1}}{D_1} \\
&+ \frac{1}{6}[W_{2c}x_{2c}(1-a_{2c}) + W_{2s}x_{2s}(1-a_{2s})]\frac{p_{CC,2}}{D_2}, \\
\overline{W}x'_{3c} =& [W_{4c}x_{4c}(1-a_{4c}) + W_{4s}x_{4s}(1-a_{4s})]p_{CI,0} \\
&+ \frac{1}{2}[W_{3c}x_{3c}(1-a_{3c}) + W_{3s}x_{3s}(1-a_{3s})]\frac{p_{CC,1} + p_{CI,1}}{D_1} \\
&+ [W_{2c}x_{2c}(1-a_{2c}) + W_{2s}x_{2s}(1-a_{2s})]\frac{\frac{1}{6}p_{CI,2} + \frac{2}{3}p_{CC,2}}{D_2} \\
&+ \frac{1}{2}[W_{1c}x_{1c}(1-a_{1c}) + W_{1s}x_{1s}(1-a_{1s})]\frac{p_{CC,3}}{D_3}, \\
\overline{W}x'_{2c} =& [W_{4c}x_{4c}(1-a_{4c}) + W_{4s}x_{4s}(1-a_{4s})]p_{II,0} \\
&+ \frac{1}{2}[W_{3c}x_{3c}(1-a_{3c}) + W_{3s}x_{3s}(1-a_{3s})]\frac{p_{II,1} + p_{CI,1}}{D_1} \\
&+ [W_{2c}x_{2c}(1-a_{2c}) + W_{2s}x_{2s}(1-a_{2s})]\frac{\frac{1}{6}p_{II,2} + \frac{2}{3}p_{CI,2} + \frac{1}{6}p_{CC,2}}{D_2} \\
&+ \frac{1}{2}[W_{1c}x_{1c}(1-a_{1c}) + W_{1s}x_{1s}(1-a_{1s})]\frac{p_{CI,3} + p_{CC,3}}{D_3} + W_0x_0\frac{p_{CC,4}}{D_4}, \\
\overline{W}x'_{1c} =& \frac{1}{2}[W_{3c}x_{3c}(1-a_{3c}) + W_{3s}x_{3s}(1-a_{3s})]\frac{p_{II,1}}{D_1} \\
&+ [W_{2c}x_{2c}(1-a_{2c}) + W_{2s}x_{2s}(1-a_{2s})]\frac{\frac{2}{3}p_{II,2} + \frac{1}{6}p_{CI,2}}{D_2} \\
&+ \frac{1}{2}[W_{1c}x_{1c}(1-a_{1c}) + W_{1s}x_{1s}(1-a_{1s})]\frac{p_{II,3} + p_{CI,3}}{D_3} + W_0x_0\frac{p_{CI,4}}{D_4}.
\end{aligned} \tag{A11}$$

$S_C$ **dominant.** In this paragraph, we consider that the SC allele is dominant over all SI alleles. In that case, all individuals that have at least one SC allele are self-compatible, and their pollen grains are compatible with all individuals in the population.

We define the frequency of gametes produced by an individual with $k$ SC alleles, with $k \in [\![1, 4]\!]$, as follows:

$$g_{k0} = \frac{1}{\overline{W}}(W_{kc}x_{kc} + W_{ks}x_{ks}). \tag{A12}$$

We also define the frequency of gametes produced by an individual with no SC alleles, in which all SI alleles are different from the $l$ given S-alleles with $l \in [\![0, 4]\!]$:

$$g_{0l} = \frac{W_0 x_0}{\overline{W}} \frac{(n-l)^4}{n^4}. \tag{A13}$$

With the above frequencies, we define the frequency of pollen with genotypes $S_C S_C$, $S_C S_i$ and $S_i S_j$ in the population, respectively, that are compatible with a maternal parent that has $k$ SI alleles:

$$\begin{aligned} r_{CC} &= g_{40} + \frac{1}{2}g_{30} + \frac{1}{6}g_{20}, \\ r_{CI} &= \frac{1}{2}g_{30} + \frac{2}{3}g_{20} + \frac{1}{2}g_{10}, \\ r_{II,k} &= \frac{1}{6}g_{20} + \frac{1}{2}g_{10} + g_{0k}. \end{aligned} \tag{A14}$$

We also define the frequency of gametes compatible with a maternal parent that has $k$ SI alleles as follows:

$$d_k = g_{40} + g_{30} + g_{20} + g_{10} + g_{0k}. \tag{A15}$$

The frequencies in the next generation are given by the following equations:

$$
\begin{aligned}
\overline{W}x'_{4s} =& W_{4c}x_{4c}a_{4c} + W_{4s}x_{4s}a_{4s} + \frac{1}{4}[W_{3c}x_{3c}a_{3c} + W_{3s}x_{3s}a_{3s}] + \frac{1}{36}[W_{2c}x_{2c}a_{2c} + W_{2s}x_{2s}a_{2s}],\\
\overline{W}x'_{3s} =& \frac{1}{2}[W_{3c}x_{3c}a_{3c} + W_{3s}x_{3s}a_{3s}] + \frac{2}{9}[W_{2c}x_{2c}a_{2c} + W_{2s}x_{2s}a_{2s}],\\
\overline{W}x'_{2s} =& \frac{1}{4}[W_{3c}x_{3c}a_{3c} + W_{3s}x_{3s}a_{3s}] + \frac{1}{2}[W_{2c}x_{2c}a_{2c} + W_{2s}x_{2s}a_{2s}] + \frac{1}{4}[W_{1c}x_{1c}a_{1c} + W_{1s}x_{1s}a_{1s}],\\
\overline{W}x'_{1s} =& \frac{2}{9}[W_{2c}x_{2c}a_{2c} + W_{2s}x_{2s}a_{2s}] + \frac{1}{2}[W_{1c}x_{1c}a_{1c} + W_{1s}x_{1s}a_{1s}],\\
\overline{W}x'_{4c} =& [W_{4c}x_{4c}(1-a_{4c}) + W_{4s}x_{4s}(1-a_{4s})]r_{CC} + \frac{1}{2}[W_{3c}x_{3c}(1-a_{3c}) + W_{3s}x_{3s}(1-a_{3s})]\frac{r_{CC}}{d_1}\\
&+ \frac{1}{6}[W_{2c}x_{2c}(1-a_{2c}) + W_{2s}x_{2s}(1-a_{2s})]\frac{r_{CC}}{d_2},\\
\overline{W}x'_{3c} =& [W_{4c}x_{4c}(1-a_{4c}) + W_{4s}x_{4s}(1-a_{4s})]r_{CI} + \frac{1}{2}[W_{3c}x_{3c}(1-a_{3c}) + W_{3s}x_{3s}(1-a_{3s})]\frac{r_{CC}+r_{CI}}{d_1}\\
&+ [W_{2c}x_{2c}(1-a_{2c}) + W_{2s}x_{2s}(1-a_{2s})]\frac{\frac{1}{6}r_{CI} + \frac{2}{3}r_{CC}}{d_2} + \frac{1}{2}[W_{1c}x_{1c}(1-a_{1c}) + W_{1s}x_{1s}(1-a_{1s})]\frac{r_{CC}}{d_3},\\
\overline{W}x'_{2c} =& [W_{4c}x_{4c}(1-a_{4c}) + W_{4s}x_{4s}(1-a_{4s})]r_{II,0} + \frac{1}{2}[W_{3c}x_{3c}(1-a_{3c}) + W_{3s}x_{3s}(1-a_{3s})]\frac{r_{II,1}+r_{CI}}{d_1}\\
&+ [W_{2c}x_{2c}(1-a_{2c}) + W_{2s}x_{2s}(1-a_{2s})]\frac{\frac{1}{6}r_{II,2} + \frac{2}{3}r_{CI} + \frac{1}{6}r_{CC}}{d_2}\\
&+ \frac{1}{2}[W_{1c}x_{1c}(1-a_{1c}) + W_{1s}x_{1s}(1-a_{1s})]\frac{r_{CI}+r_{CC}}{d_3} + W_0x_0\frac{r_{CC}}{d_4},\\
\overline{W}x'_{1c} =& \frac{1}{2}[W_{3c}x_{3c}(1-a_{3c}) + W_{3s}x_{3s}(1-a_{3s})]\frac{r_{II,1}}{d_1} + [W_{2c}x_{2c}(1-a_{2c}) + W_{2s}x_{2s}(1-a_{2s})]\frac{\frac{2}{3}r_{II,2} + \frac{1}{6}r_{CI}}{d_2}\\
&+ \frac{1}{2}[W_{1c}x_{1c}(1-a_{1c}) + W_{1s}x_{1s}(1-a_{1s})]\frac{r_{II,3}+r_{CI}}{d_3} + W_0x_0\frac{r_{CI}}{d_4}.
\end{aligned}
\tag{A16}
$$

##### A.1.3 Diploids: Dominance classes

In this model, SI alleles are distributed among dominance classes, while the SC allele either belongs to one of these classes or is dominant over all SI alleles. Unlike the previous model, the population is structured according to ordered S-genotypes, denoted $S_iS_j$ where $S_i$ and $S_j$ can also be $S_C$, rather than into groups. Consequently, each possible genotype is explicitly considered in the model. The frequencies of individuals with genotype $S_iS_j$ that come from selfing and outcrossing at the next generation can be expressed as:

$$\overline{W}x^{\text{c'}}_{ij} = \sum_{M} \left[ [W^{\text{c}} x^{\text{c}}_{M}(1 - a^{\text{c}}_{M}) + W^{\text{s}} x^{\text{s}}_{M}(1 - a^{\text{s}}_{M})] \times \frac{\sum_F \mathrm{C}_{MF} \mathrm{P}_{MF\to ij}(W^{\text{c}} x^{\text{c}}_{F} + W^{\text{s}} x^{\text{s}}_{F})}{\sum_F \mathrm{C}_{MF}(W^{\text{c}} x^{\text{c}}_{F} + W^{\text{s}} x^{\text{s}}_{F})} \right]$$

$$\overline{W}x^{\text{s'}}_{ij} = \sum_{M} [W^{\text{c}} x^{\text{c}}_{M} a^{\text{c}}_{M} + W^{\text{s}} x^{\text{s}}_{M} a^{\text{s}}_{M}] \times \mathrm{C}_{MM} \tag{A17}$$

where $M = (i_M, j_M)$ and $F = (i_F, j_F)$ are the S-genotypes of the mother and father respectively. The function $\mathrm{C}_{MF}$ gives the compatibility between individual $M$ and $F$, i.e. $\mathrm{C}_{MF} = 1$ if individuals $M$ and $F$ are compatible, and 0 if they are not. The function $\mathrm{P}_{MF\to ij}$ gives the probability for individuals $M$ and $F$ to form an offspring with genotypes $S_iS_j$.

Rather than deriving an explicit recursion equation for each genotype, these expressions are evaluated algorithmically in *Julia*. At each generation, the algorithm iterates over all possible maternal genotypes $M$. For each mother, all compatible paternal genotypes $F$ are considered, and their contributions are weighted by their frequency, fitness, and the probability of producing offspring with genotype $S_iS_j$. These contributions are summed over all compatible fathers and normalized by the total frequency of compatible pollen to account for pollen competition. Finally, the contributions of all maternal genotypes are summed to obtain the genotype frequencies in the next generation.

#### A.2 Analytical resolutions

This section of the appendix provides a detailed analytical resolution of the models presented in Sections 2.1.1 and 2.1.2. For both diploids and tetraploids, the resolution was carried out using *Wolfram Mathematica* (notebooks available) and consists of the following steps:

1. Linearize the recursion equations to first order.
2. Build a matrix $J$, such that $X' = JX$, where $X$ is the vector of frequencies.
3. Evaluate the Routh-Hurwitz stability conditions using the characteristic polynomial of $J$.

When the Routh-Hurwitz conditions are satisfied, the absence of SC alleles is a stable equilibrium, indicating complete maintenance of the SSI system.

##### A.2.1 Diploids: SI codominant

**$S_C$ codominant or recessive.** After linearizing the recursions equations (A2) to first order, the system simplifies to a $2 \times 2$ matrix:

$$\begin{pmatrix} x_1' \\ x_4' \end{pmatrix} = \begin{pmatrix} \frac{\alpha(\delta-1)^2}{1-\alpha\delta} & 0 \\ -\frac{(\delta-1)\left(2(\alpha-1)+n\left(4-4\alpha+n(-2+\alpha+\alpha\delta)\right)\right)}{(2+(-4+n)n)(-1+\alpha\delta)} & 1+\frac{n-1}{2+(-4+n)n} \end{pmatrix} \begin{pmatrix} x_1 \\ x_4 \end{pmatrix} \tag{A18}$$

The frequency $x_2$ does not appear in this equation, as the frequency of $S_C S_C$ individuals produced by outcrossing becomes negligible when the SC allele is rare in the population.

For a $2 \times 2$ matrix, the characteristic polynomial is a quadratic equation of the form:

$$r^2 + c_1 r + c_2 = 0 \tag{A19}$$

with:

$$\begin{aligned} c_1 &= 1 + \frac{1}{2+(-4+n)n} - \frac{n}{2+(-4+n)n} + \frac{\alpha(\delta-1)^2}{-1+\alpha\delta} \\ c_2 &= -\frac{(n-1)\left(-1+\alpha+\alpha(\delta-1)\delta\right)}{(2+(-4+n)n)(-1+\alpha\delta)} \end{aligned} \tag{A20}$$

The coefficient $c_2$ is always negative for realistic numbers of S-alleles in the population, which prevents the Routh-Hurwitz conditions from being satisfied. This indicates that the equilibrium where SC is absent is never stable. In other words, the SC allele is always present in the population at equilibrium: it either invades the population or coexists with SI alleles.

**$S_C$ dominant.** When the SC allele is dominant over all SI alleles, the recursion equations (A3) to first order can be rewritten as a $3 \times 3$ matrix:

$$\begin{pmatrix} x_1' \\ x_3' \\ x_4' \end{pmatrix} = \begin{pmatrix} -\frac{\alpha(\delta-1)^2}{-1+\alpha\delta} & -\frac{\alpha(\delta-1)^2}{-4+4\alpha\delta} & \frac{\alpha}{4} \\ 0 & -\frac{\alpha(\delta-1)^2}{-2+2\alpha\delta} & \frac{\alpha}{2} \\ j_{31} & j_{32} & \frac{(n-1)^2}{2+(-4+n)n} - \frac{\alpha}{2} \end{pmatrix} \begin{pmatrix} x_1 \\ x_3 \\ x_4 \end{pmatrix} \tag{A21}$$

with:

$$
\begin{aligned}
j_{31} &= -\frac{(\delta-1)\left(2(\alpha-1)+n\left(4-4\alpha+n(-2+\alpha+\alpha\delta)\right)\right)}{(2+(-4+n)n)(-1+\alpha\delta)} \\
j_{32} &= -\frac{(\delta-1)\left(2(\alpha-1)+n\left(4-4\alpha+n(-2+\alpha+\alpha\delta)\right)\right)}{2(2+(-4+n)n)(-1+\alpha\delta)}
\end{aligned}
\tag{A22}
$$

The corresponding characteristic polynomial is of the form:

$$
r^3 + c_1 r^2 + c_2 r + c_3 = 0 \tag{A23}
$$

Two of the Routh-Hurwitz conditions ($c_1 > 0$ and $c_1 c_2 > 0$) seem to be verified for all sets of realistic parameters $(\alpha, \delta, n)$, but the coefficient $c_3$ can change sign depending on the parameter values. We have:

$$
c_3 = \frac{\left(-2+\alpha+\alpha\delta^2\right)\Big(2(\alpha-1)+n\big(4+(-4+n)\alpha\big)-\alpha\big(4+n(-8+n(2+\alpha))\big)\delta+2\alpha\big(1-2n+(-1+n)^2\alpha\big)\delta^2\Big)}{4\left(2+(-4+n)n\right)\left(-1+\alpha\delta\right)^2} \tag{A24}
$$

Considering $c_3$ as a polynomial in $\delta$, we solve the equation $c_3 = 0$ to obtain the threshold inbreeding depression necessary to prevent the invasion of SC allele (Figure 2).

##### A.2.2 Tetraploids: SI codominant

**$S_C$ codominant or recessive.** After linearization to first order, the recursion equations (A11) become:

$$
\begin{pmatrix} x'_{4s} \\ x'_{1c} \\ x'_{2c} \end{pmatrix} = \begin{pmatrix} -\dfrac{\alpha(\delta-1)^2}{-1+\alpha\delta} & 0 & 0 \\ 0 & 1+\dfrac{2}{-7+n} & \dfrac{4}{3}+\dfrac{40}{3(-7+n)}-\dfrac{8}{-6+n} \\ j_{31} & 0 & \dfrac{1}{3}+\dfrac{10}{3(-7+n)}-\dfrac{2}{-6+n} \end{pmatrix} \begin{pmatrix} x_{4s} \\ x_{1c} \\ x_{2c} \end{pmatrix} \tag{A25}
$$

with:

$$
\begin{aligned}
j_{31} =& \frac{(1-\delta)}{(-7+n)(-6+n)(-5+n)(-4+n)(-1+\alpha\delta)} \times [840(\alpha-1) \\
&+ n\big(644-2n(95+(-14+n)n)-638\alpha+n(179+(-22+n)n)\alpha+(-3+n)(-2+n)(-1+n)\alpha\delta\big)]
\end{aligned}
\tag{A26}
$$

The characteristic polynomial is of the form of equation (A23), where:

$$c_3 = -\frac{4\,(51 + (-15+n)n)\,(-1+\alpha+\alpha(\delta-1)\delta)}{3\,(-7+n)^2\,(-6+n)\,(-1+\alpha\delta)} \tag{A27}$$

The coefficient $c_3$ is always negative for positive parameter values of $(\alpha, \delta, n)$, indicating that the SC allele is always present in the population.

**$S_C$ dominant.** In this case, the recursion equations (A16) can be rewritten as a $6 \times 6$ matrix. The Routh-Hurwitz conditions require that the determinants of matrices constructed using the coefficients of the characteristic polynomial are all positive. We were unable to determine analytically the parameter sets that verify the positivity of all these determinants. Therefore, this case was only solved numerically by iterating the recursion equations until an equilibrium was reached (Figure 3).

### B Supplementary Figures

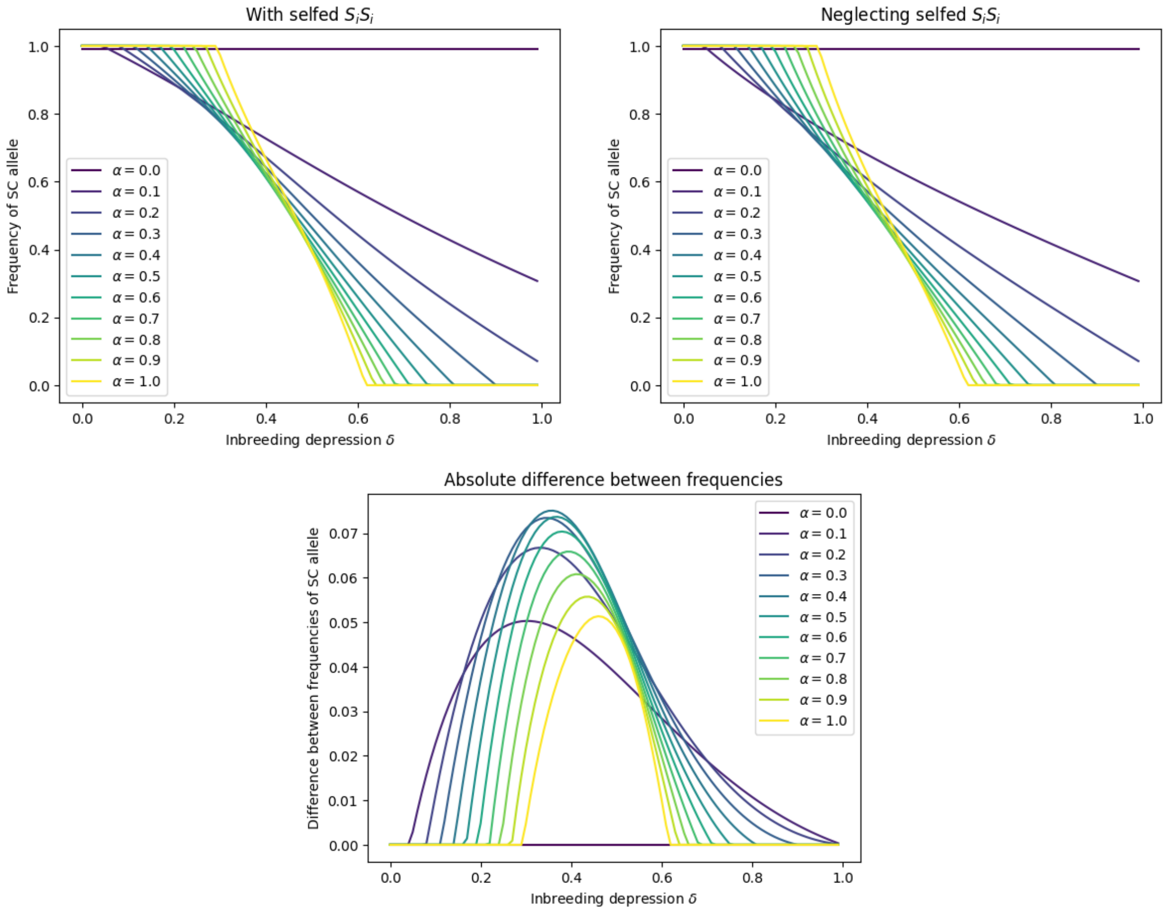


Figure S1: Comparison between the results obtained using two versions of the analytical model for diploids when the SC allele is dominant by computing the absolute value of the difference between the frequencies of SC alleles at equilibrium (bottom): one model includes the fitness difference in selfed $S_iS_i$ individuals that suffer from inbreeding depression (exact model: top left), and the other neglects this fitness difference (approximate model: top right).

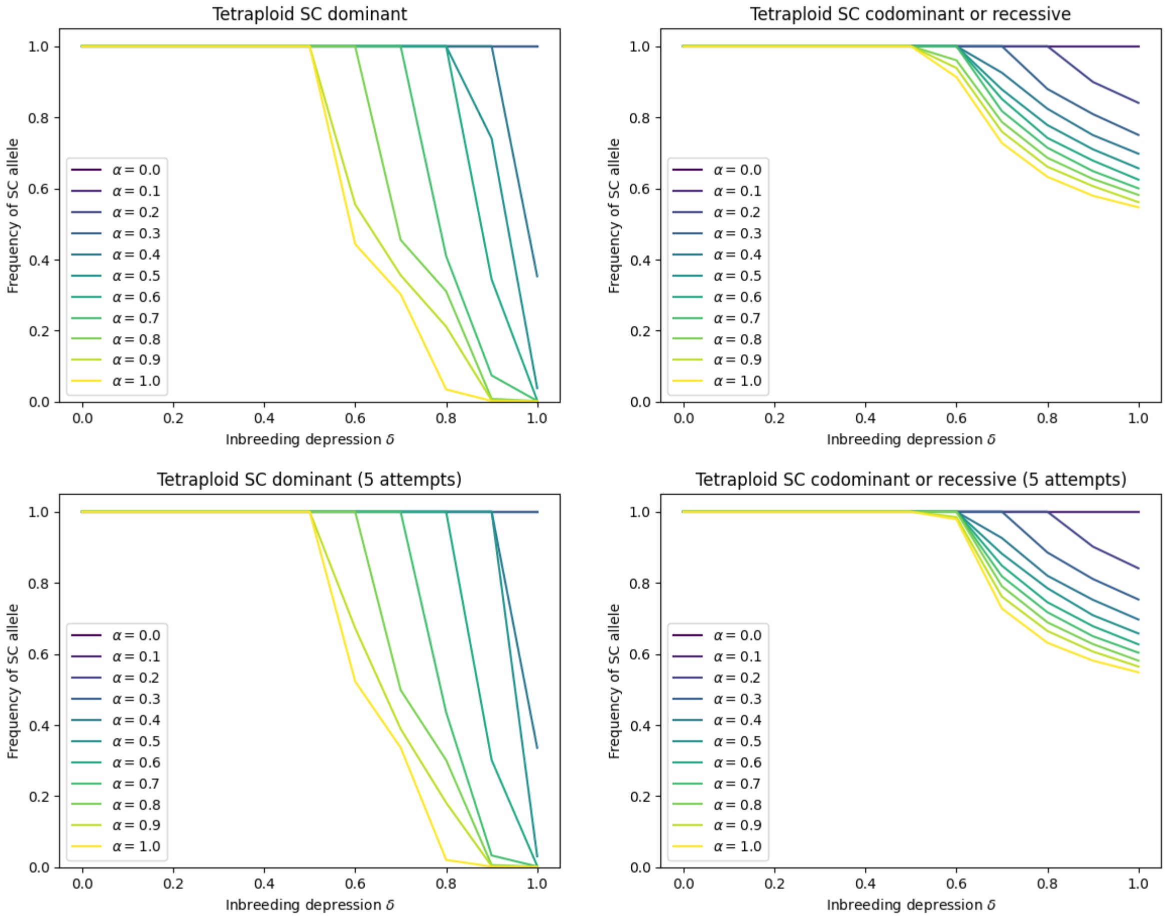


Figure S2: Comparison between two different pollen limitation scenarios in tetraploids when SI alleles are codominant. Frequency of SC allele at equilibrium as a function of inbreeding depression for different values of the self-pollen rate (represented by different colored lines) when $N_{\mathrm{pop}} = 1000$. Results obtained using the individual-based model for tetraploids where inbreeding depression $\delta$ is fixed when the mother is rejected after 50 pollination attempts (top) and when the mother is rejected after 5 attempts only (bottom) when the SC allele is dominant (left) or codominant/recessive (right).

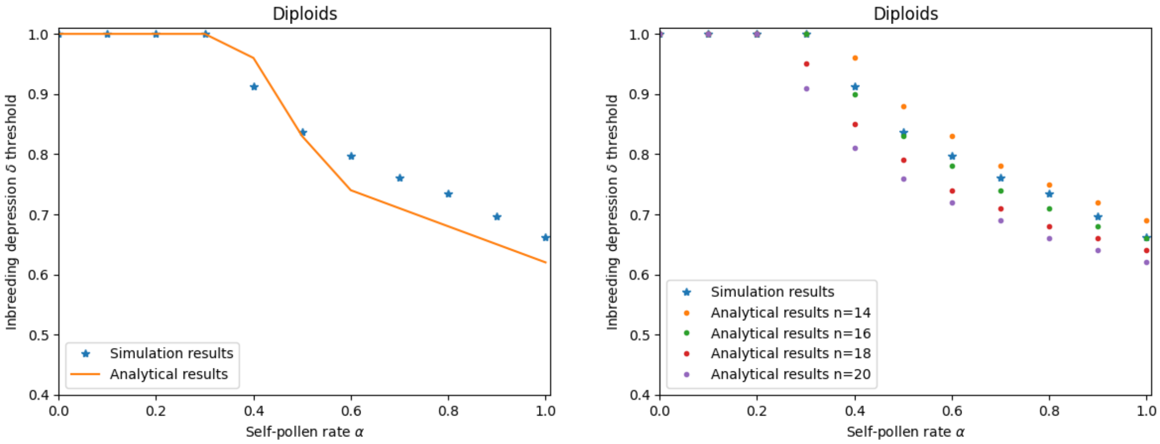


Figure S3: SC allele dominant over all SI alleles, when SI alleles are codominant in a diploid population. Minimal inbreeding depression necessary to prevent the maintenance of an SC mutant allele into an initially SI population obtained using individual-based simulations (5 runs) where inbreeding depression is controlled by the rate of deleterious mutations $U$. Comparison between simulation results and analytical results with the corresponding number of SI alleles between 13 and 20 depending on the self-pollen rate $\alpha$ (on the left), or with different values of the number of SI alleles (on the right).

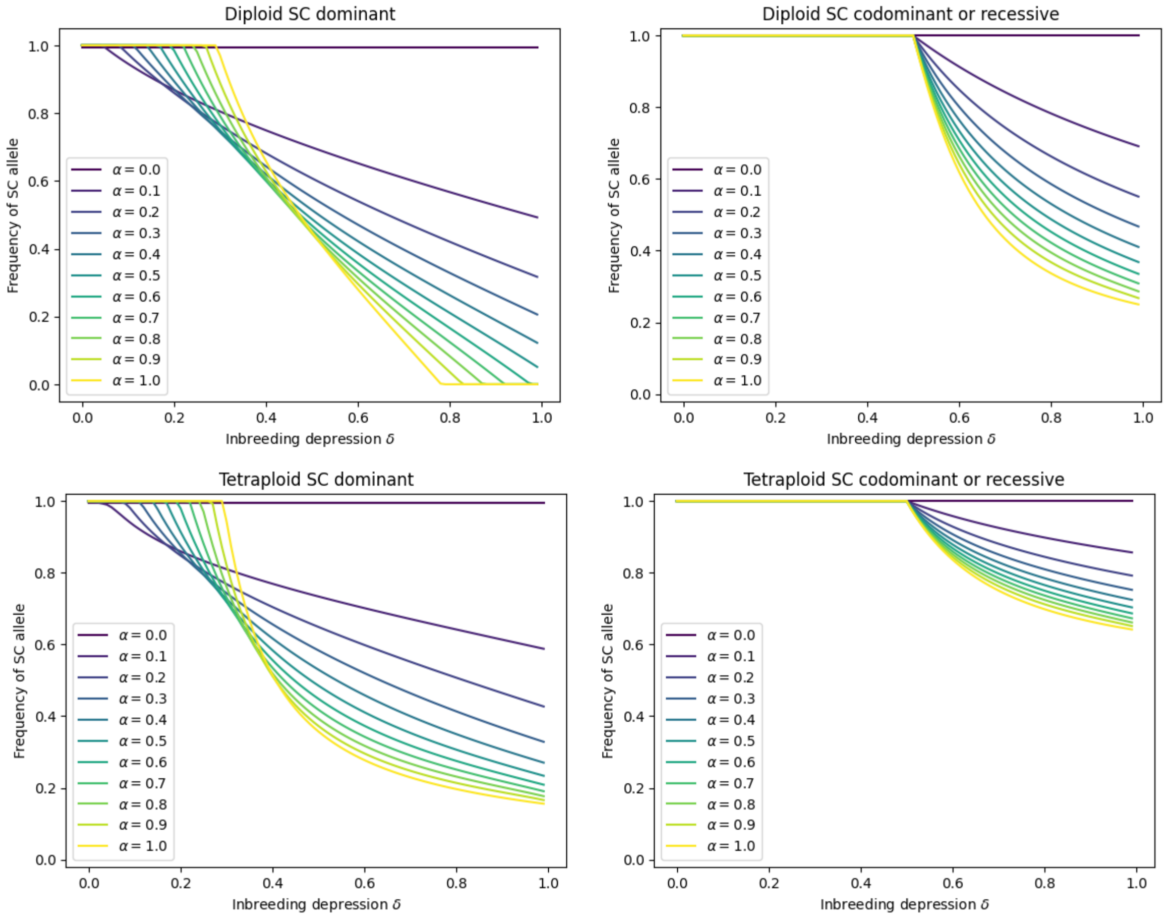


Figure S4: Frequency of SC allele at equilibrium as a function of inbreeding depression for different values of the self-pollen rate (represented by different colored lines) when $n = 10$. Results obtained by recursions on the analytical models for diploids (top) and for tetraploids (bottom) when the SC allele is dominant (left) or codominant/recessive (right).

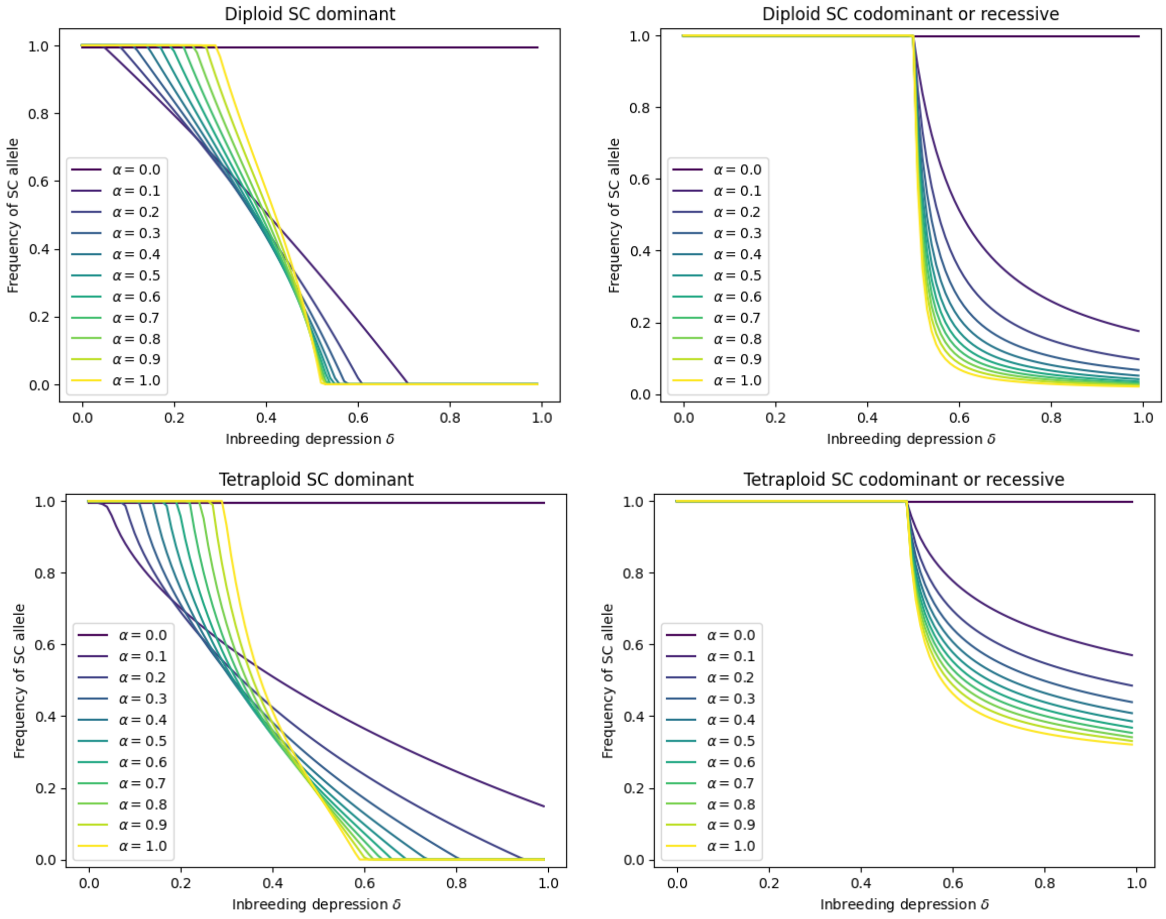


Figure S5: Frequency of SC allele at equilibrium as a function of inbreeding depression for different values of the self-pollen rate (represented by different colored lines) when $n = 100$. Results obtained by recursions on the analytical models for diploids (top) and for tetraploids (bottom) when the SC allele is dominant (left) or codominant/recessive (right).

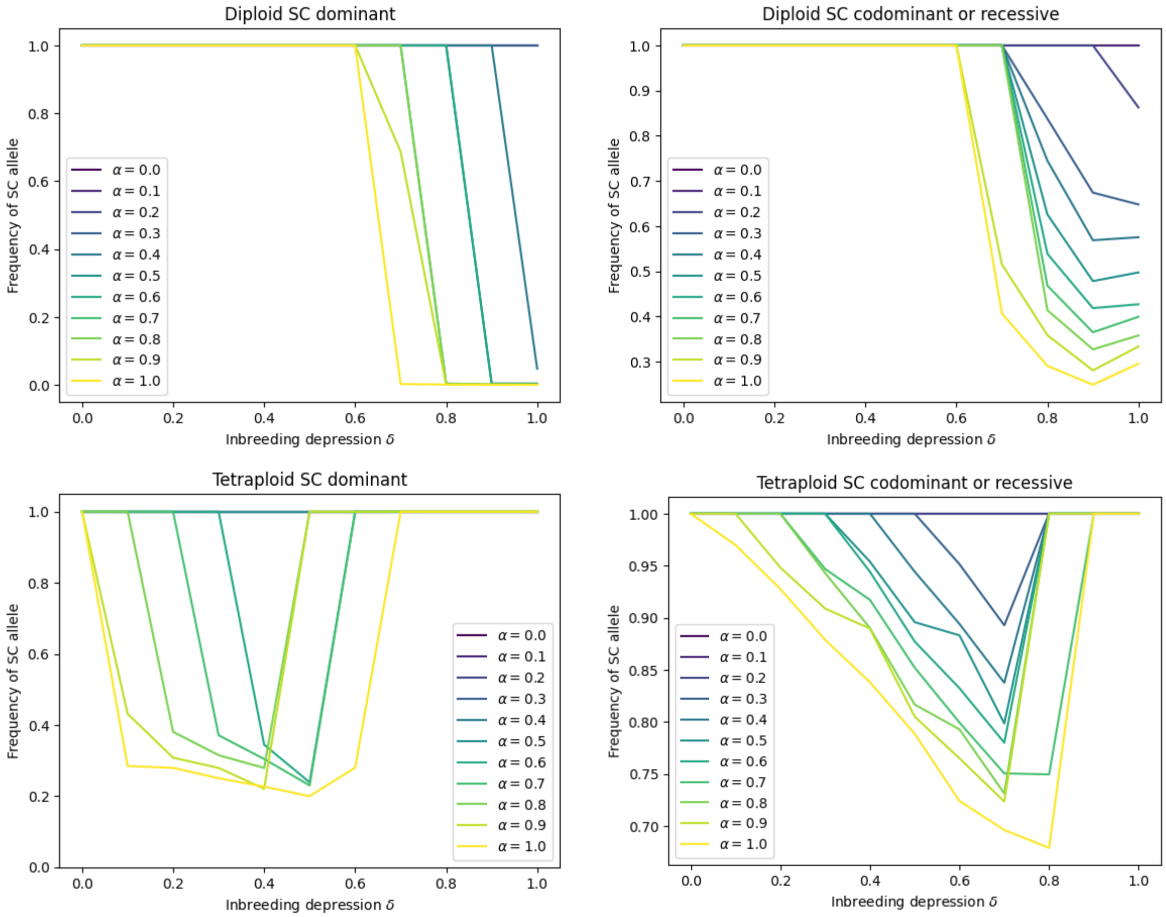


Figure S6: Frequency of SC allele at equilibrium as a function of inbreeding depression for different values of the self-pollen rate (represented by different colored lines) when $N_{\text{pop}} = 1000$. Results obtained using the individual-based model where inbreeding depression $\delta$ is determined by the rate of deleterious mutations $U$ for diploids (top) and tetraploids (bottom) when the SC allele is dominant (left) or codominant/recessive (right).
